# Defining bottlenecks and physiological impact of an orthogonal translation initiation system

**DOI:** 10.64898/2025.12.21.695813

**Authors:** Dominic Scopelliti, Russel M. Vincent, Andras Hutvagner, Fiona Whelan, William P. Klare, Ignatius Pang, George M. Church, Paul R. Jaschke

**Author notes:** Correspondence to Paul R Jaschke. These authors contributed equally (Dominic Scopelliti, Russel M. Vincent, Andras Hutvagner).

## Abstract

Reengineering translation initiation provides a powerful route to develop new translation systems that enable precise control of protein synthesis. While many engineered translation systems show promise, their orthogonality and impact on host physiology is largely uncharacterized, limiting broader application. Here, we develop an initiator-tRNA with an AAC anticodon mutation (i-tRNA-AAC), enabling translation initiation at a GUU start codon. Using fluorescence assays, proteomics, and tRNA sequencing, we assess the i-tRNA-AAC initiation orthogonality and effects on the host, guiding the optimization of its translational efficiency. We find the i-tRNA-AAC mutant initiates translation exclusively from its GUU start codon and is improved by overexpression of valyl-tRNA synthetase and methionyl-tRNA formyltransferase. However, this intervention perturbs aminoacylation and base modifications of endogenous tRNAs along with proteome-wide changes likely due to increased valine demand. Our findings demonstrate how an orthogonal translation initiation system reshapes host physiology and reveals the adaptive responses that accompany translational reprogramming.

## Introduction

Orthogonal translation systems (OTSs), comprised of heterologous translation components that function independently of the host’s native machinery, have emerged as a powerful tool in the field of synthetic biology. These systems have enabled genetic code expansion and the precise reprogramming of protein synthesis^1–6^. Through the reengineering and decoupling of translational machinery from their endogenous counterparts, OTSs have facilitated enhanced control over protein expression, the development of insulated synthetic circuits, and the incorporation of non-canonical amino acids into proteins^7–18^. However, while most previous research efforts have focused on reengineering internal codons, there has been increasing interest in targeting the initiation phase of translation through the start codon^19–23^.

In the standard genetic code, the AUG triplet encodes methionine and is used both in translation initiation and at internal positions of a coding sequence. These dual roles of AUG are distinguished by two tRNA species: the initiator tRNA (i-tRNA), which recognizes AUG at the translation initiation region, and methionyl elongator tRNAs (e-tRNAs), which decode internal AUG codons. Although both tRNAs share the same anticodon and are aminoacylated by methionyl-tRNA synthetase, the i-tRNA carries distinct sequence and structural features. They are, the C1 x A72 mismatch, A11-U24 base-pair, and 3GC pairs in the anticodon stem which enable its recruitment by initiation factors, direct binding to the ribosomal P-site, and ribosome maturation^24–28^. In bacteria, only i-tRNAs charged with methionine are formylated^29^, selectively increasing i-tRNA affinity for initiation factor 2 and promoting recruitment to the 30S ribosomal pre-initiation complex^26^. Because translation initiation occurs in a defined positional context and uses specialized machinery distinct from elongation, it provides a unique opportunity to add context-specific control to synthetic circuits and to incorporate non-canonical amino acids at the N-terminus of proteins^19,20^.

Mutations in the anticodon of i-tRNAs are known to change which codons are capable of initiating translation as well as which amino acid is incorporated onto the N-terminus of the target protein^19–22,30–32^. The amino acid that a tRNA carries is defined by its molecular interactions with tRNA-aminoacyl synthetases that selectively charge tRNAs with an amino acid, thereby directly maintaining the fidelity of the genetic code. Aminoacyl synthetases (aaRSs) have a range of positive and negative tRNA sequence determinants that modulate relative amino acid charging efficiency between similar tRNA substrates^33^. Understanding the physiological consequences^22,34,35^ and molecular details of how tRNA anticodon mutants with hybrid aaRS sequence determinants interact with aminoacyl synthetases and the translation machinery has ramifications not only for the success of genetic code expansion efforts^5,7,9–11^ but also characterizing and treating human diseases^36^.

Here, we use multi-omics to define, at an unprecedented depth, the *in vivo* effects of deploying an OTS and overexpression of translation-associated enzymes. We show that this i-tRNA-AAC mutant initiates translation completely orthogonally from its cognate GUU start codon and predominantly incorporates valine at protein N-termini. Using mass spectrometry, structural modelling, and tRNA profiling strategies, we identify tRNA maturation bottlenecks and assess changes in the global proteome in response to the expression of the i-tRNA mutant and heterologous translation associated enzymes. By quantifying system orthogonality, identifying rate limiting steps, and mapping host responses, these results provide a framework to balance yield, fidelity, and host compatibility in future i-tRNA-based OTSs.

## Results

### Mutant i-tRNA-AAC selectively initiates translation from the non-canonical GUU start codon with valine and its formylated derivative

To evaluate the translational competence and orthogonality of an i-tRNA bearing a mutated anticodon, we measured its capacity to initiate translation from all 64 potential start codons. To achieve this, we began with the *E. coli metY* gene encoding i-tRNA-fMet-CAU-2 and mutated the anticodon to AAC, hereafter referred to as i-tRNA-AAC (Figure S1). This construct was cloned into the IPTG-inducible pULTRA*-tac* vector, and a negative control plasmid lacking the *metY* gene was built in parallel. We then co-transformed either the pULTRA-*tac*-*metY(AAC)* or the empty control vector with a previously described library of sfGFP reporters controlled by each of the 64 possible start codons (pET20b-*T7-(NNN)sfGFP*, where NNN denotes the start codon) (Figure 1A)^37^.

**Figure 1.**
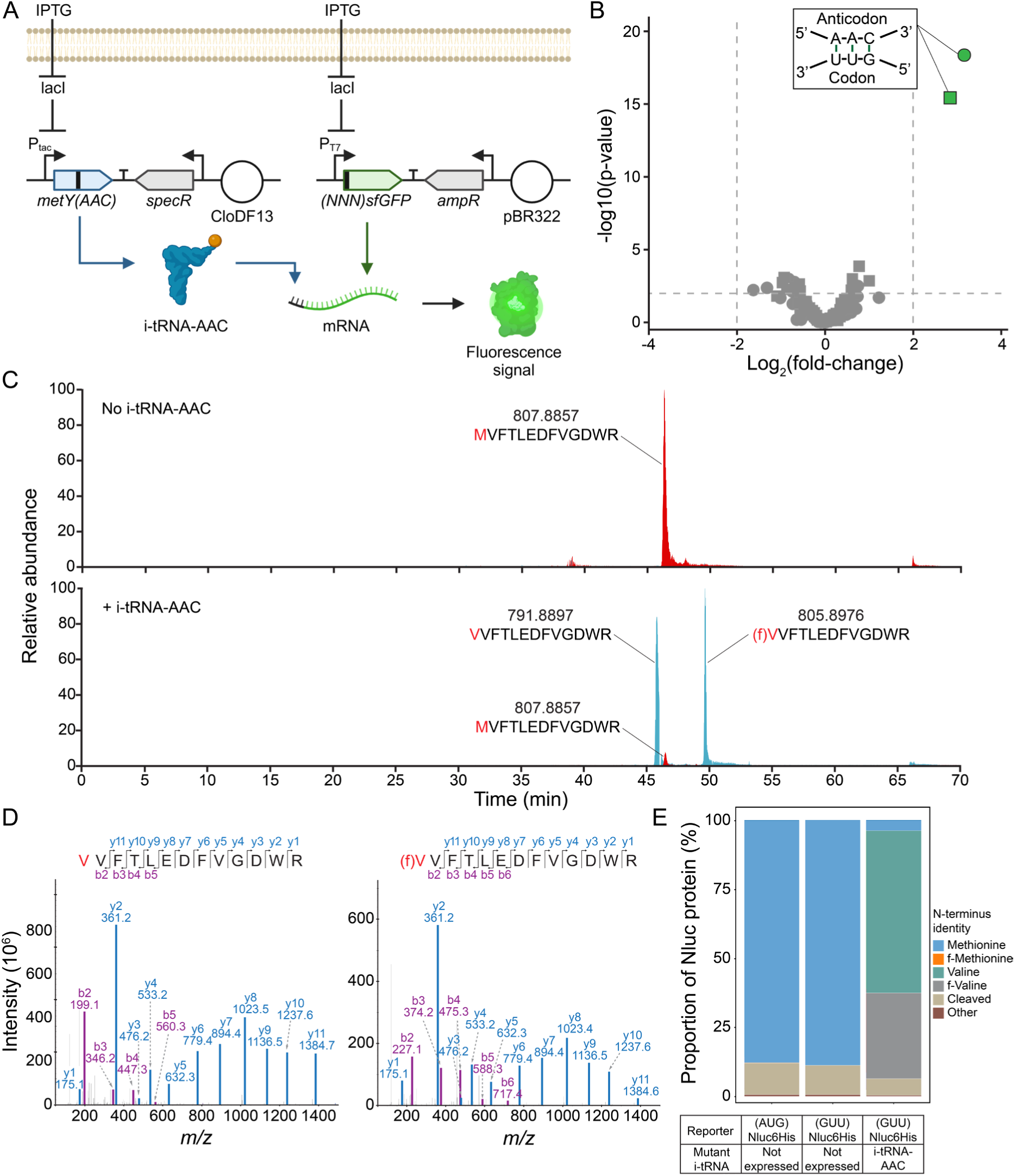
Mutant i-tRNA-AAC selectively initiates translation from the non-canonical GUU start codon with valine and its formylated derivative. (A) Schematic overview of the i-tRNA-AAC mediated translation initiation system. The mutant i-tRNA-AAC was expressed from an IPTG-inducible tac promoter on a pULTRA plasmid. Co-expressed is each variant from a 64-start codon library encoding a (NNN)sfGFP reporter from the T7 promoter on a pET20B vector. Created in BioRender. J, P. (2025) https://BioRender.com/44leygd (B) DEseq2 analysis comparing the translation efficiencies between strains expressing the i-tRNA-AAC and empty vector control, using the pET20b-*T7-(NNN)sfGFP* reporter library. A significant increase in fluorescence from GUU start codons was observed in the presence of i-tRNA-AAC (green), while all other start codons remain unaffected (grey). Bulk fluorescence data are indicated by squares while flow cytometry data shown by circles. (C) Extracted ion chromatograms (XICs) of MS1 data from N-terminal tryptic peptides in the presence and absence of i-tRNA-AAC. (D) MS/MS fragmentation spectra of the N-terminal VVFTLEDFVGDWR (m/z 791.8897) and (f)VVFTLEDFVGDWR (m/z 805.8976) confirm the presence of valine and formylated valine-initiated (GUU)Nluc proteins. Unique b-type fragment ions were used for positive identification. (E) Relative abundance of N-terminal peptides initiated with different amino acids from (AUG)Nluc and (GUU)Nluc reporter proteins.

Fluorescence intensity at both the population and single-cell level served as a quantitative readout of translation efficiency. As expected from previous work^37^, in the pULTRA-empty control we observed unimodal translation initiation from multiple non-cognate start codons (Figures S2 and S3). By contrast, in strains carrying pULTRA-*tac*-*metY(AAC)* we observed significantly higher fluorescence from the reporter carrying the cognate GUU start codon (Figure S2). Repurposing DEseq2^38^ to identify significantly upregulated translation initiation across the reporter library, we found only the cognate (GUU)sfGFP reporter satisfied these conditions, with an 8-fold increase in population level fluorescence and a 10-fold increase at the single cell level with unimodal distribution relative to the control (p < 0.01) (Figure 1B). These results indicate that the mutant i-tRNA-AAC exhibits high fidelity for its complementary GUU start codon and is functionally orthogonal with respect to all other potential start codons. This specificity suggests that engineered i-tRNAs, such as i-tRNA-AAC, could offer tighter translational control than existing systems that rely on the canonical i-tRNA-fMet-CAU and thus may serve as a valuable tool in the deployment of synthetic circuits requiring precise initiation^39,40^.

Having established the efficiency and orthogonality of translation initiation by the i-tRNA-AAC and GUU start codon pair, we next sought to determine which amino acid was incorporated at the N-terminus of proteins expressed using this system. To this end, we constructed two C-terminally His-tagged NanoLuc luciferase (Nluc) reporter plasmids bearing either a GUU or AUG start codon, namely pET20b-*T7-(GUU)Nluc6HIS* and pET20b-*T7-(AUG)Nluc6HIS*. The (NNN)Nluc reporter protein was specifically selected due to its favorable tryptic cleavage profile, which facilitates downstream analysis of N-terminal peptides using mass spectrometry. Each Nluc reporter was co-expressed with either the i-tRNA-AAC encoding plasmid (pULTRA-*tac*-*metY(AAC)*), or the empty vector control, and purified. To identify the amino acid incorporated onto the N-terminus, we employed a data-dependent acquisition (DDA) mass spectrometry workflow, which enabled us to agnostically survey for all potential N-terminal amino acids, including both their formylated and unformylated states. An inclusion list was used to prioritize the fragmentation of key peptides, including multiple internal Nluc tryptic peptides and all 20 amino acids as potential N-terminal residues (Supplementary File S1). Given that peptides would likely differ only at their N-terminal residue, the presence of unique b–type product ions were used to identify the initiating amino acid while precursor ion intensity was used for quantitation.

As expected, in control strains co-expressing pET20b-*T7-(AUG)Nluc6His* and pULTRA-*tac*-*Empty* vectors, N-terminal peptides of purified (AUG)Nluc proteins predominantly carried methionine (87.9% ± 0.5%) (Figure 1C and E), formylated methionine (0.10% ± 0.02%), or were cleaved and lacked the N-terminal amino acid (11.4% ± 0.5%) (Figure S4). Unexpectedly, we also detected low abundance peptides initiated with glycine (0.30% ± 0.02%) or threonine (0.30% ± 0.01%) (Figure S5). In strains carrying the (GUU)Nluc reporter and empty control vector, N-terminal peptides were also either lacking the N-terminal residue (11% ± 2%) or initiated using methionine (89% ± 2%) (Figure S6), consistent with prior reports of low-level initiation from non-canonical start codons by the native i-tRNA-fMet-CAU^37^.

By contrast, in strains expressing i-tRNA-AAC and the (GUU)Nluc reporter, mass spectrometry revealed that valine (59% ± 9%) was the predominant N-terminal residue (Figure 1C-E). This observation reinforces previous findings suggesting that the amino acid charged to a tRNA is primarily influenced by its anticodon sequence^22,31,32,41^. In *E. coli*, valyl-tRNA synthetase (ValRS) recognizes and aminoacylates valine isoacceptors, and likely charges i-tRNA-AAC due to the presence of known identity elements in its structure, including A35, C36, and the conserved A73 (Figure S1)^33,42,43^. In addition to valine, we also detected its formylated derivative, formyl-valine (30% ± 10%) (Figure 1C and 1D), suggesting i-tRNA-AAC charged with valine is a substrate for endogenous methionyl-tRNA formyltransferase (FMT). In addition to the valine-initiated peptides, we detected a subset of N-terminally cleaved peptides (6% ± 2%) (Figure S7A), as well as low abundance peptides bearing methionine (3.6% ± 0.2%) (Figure S7B), glycine (0.10% ± 0.03%), alanine (0.04% ± 0.01%), and isoleucine (0.14% ± 0.07%) at the N-terminus (Figure S8). The cleaved fraction likely resulted from the co-translational activity of methionine aminopeptidase (MAP)^44^. The absolute specificity of MAP for methionine residues at the N-terminus of proteins has been previously established^45^, thus we inferred that the cleaved fractions found within this study likely originated from methionine-initiated proteins. Further, we postulate that the N-terminal methionine is most plausibly explained by low-level background initiation from the native i-tRNA-fMet-CAU, as GUU start codons are weakly recognized by this initiator species^37,46^. Supporting this, methionine-initiated proteins were observed both in the presence (Figure S7B) and absence (Figure S6B) of i-tRNA-AAC expression. These findings suggest that rather than misacylation of i-tRNA-AAC with methionine, background activity from endogenous i-tRNA-fMet-CAU constitutes a low-frequency competing initiation modality.

### *In silico* analysis of formylated-valine peptide indicates inefficient interactions with peptide deformylase

Typically, in *E. coli*, methionine is charged onto i-tRNA-fMet-CAU, followed by formylation, before it can participate in translation initiation, but formylated-methionine peptides are difficult to observe due to the rapid co-translational removal of the formyl group by peptide deformylase^47^. The elevated proportion of formylated valine (30% ± 10%) in (GUU)Nluc samples compared to the proportion of formylated methionine (0.10% ± 0.02%) in (AUG)Nluc samples (Figure 1E) led us to investigate the reason for this discrepancy. We hypothesized that the same order of charging followed by formylation occurred for the i-tRNA-AAC, but that peptide deformylase was less efficient at removing the formyl group, thus leading to a large pool of proteins bearing formylated-valine (Figure 1E).

To simulate the interaction between peptide deformylase and formylated valine, we docked models of formylated peptides fMAS, fVAS and fAAS into the Ni-bound active site (1bs6, chain A). We found no profound differences in the active site localization of the formyl group in the fVAS versus fMAS peptides (Figure 2A). The fMAS peptide having the formyl carbonyl carbon coordinated in proximity to the catalytic nucleophile W1 (3.0 Å), the backbone amide hydrogen of L91 (2.3 Å), and the active site Ni atom (3.4 Å) (Figure 2B), consistent with the mechanism of peptide deformylase formyl hydrolysis proposed by Becker et al.^48^ and modelled by Wu et al.^49^. The highest scoring docked fVAS peptide showed a similar orientation, with the formyl carbonyl carbon more closely associated with W1 (2.9 Å) and Ni (3.0 Å), and slightly further from L91 (2.7 Å), likely indicating that this binding event would be competent for catalysis, but is expected to be less efficient based on the decreased accessible surface area for the side chain at position 1 (Figure 2C and 2E), and decreased hydrophobic interaction strength of the fVAS substrate in the binding pocket (Table 1).

**Figure 2.**
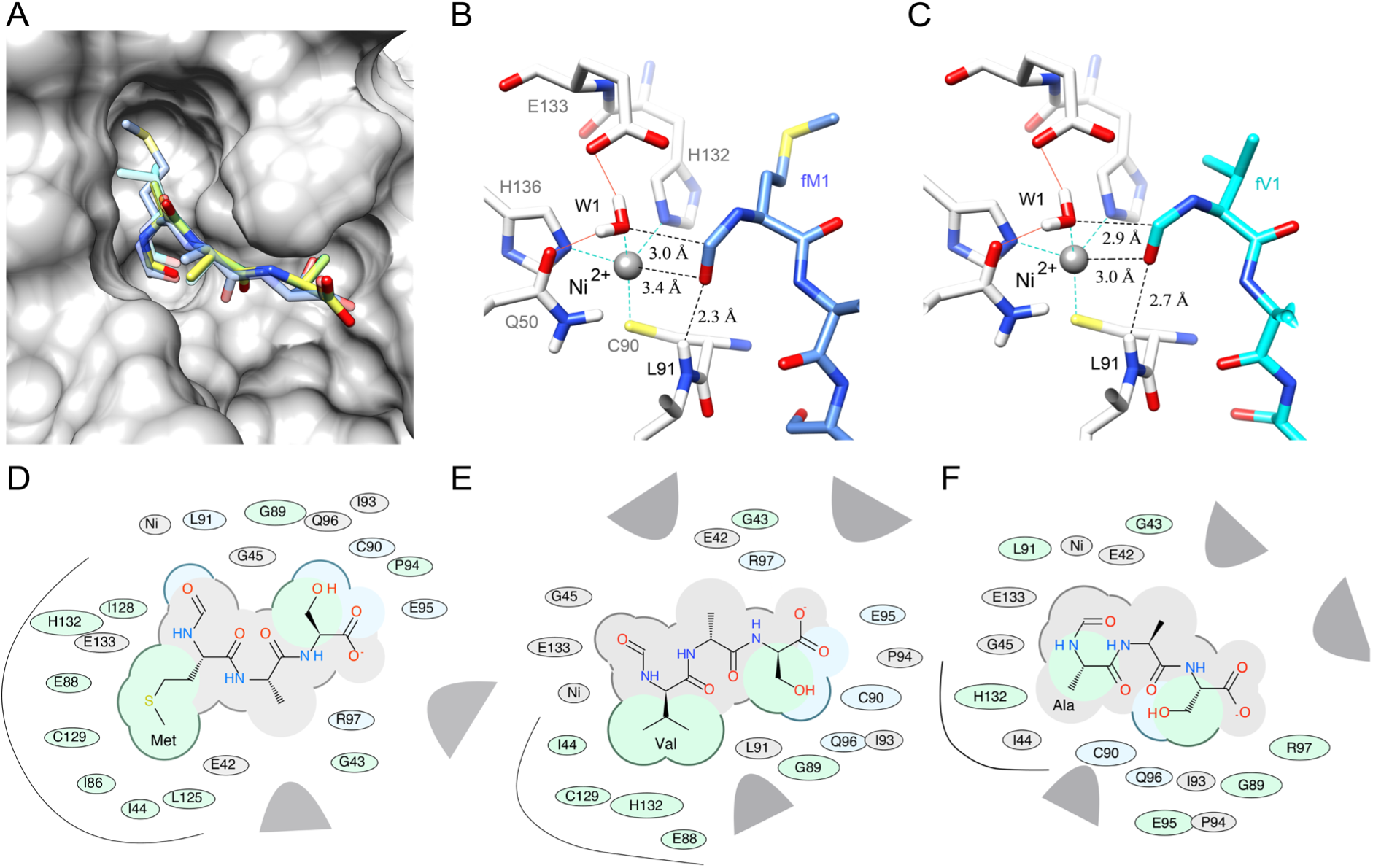
Overlay of models of highest scoring docking pose of formylated peptides fMAS, fVAS and fAAS in the substrate binding pocket of Ni^2+^ bound *E. coli* peptide deformylase. (A) A model of *E. coli* peptide deformylase (pdb code: 1bs6) is shown in grey ribbon, with the surface rendered to illustrate the peptide binding pocket (sidechains, grey sticks). The highest scoring docking poses for peptides fMAS (blue), fVAS (cyan) and fAAS (yellow) are illustrated as sticks, with atoms colored by type (sulphur, yellow; oxygen, red; nitrogen, blue). (B) The active site binding interactions of the highest scoring pose of fMAS and (C) fVAS peptides, illustrating Gln50 and Glu133 hydrogen bonding to water (W1) (red lines); and nickel (Ni^2+^, grey sphere) coordination by Cys90, His132 and His136 (cyan dashed lines). Distances between modelled formyl carbon and W1, Ni^2+^ and Leu91 backbone amide are illustrated as black dashed lines. (D) 2D chemical interaction diagram for best scoring docked fMAS, (E) fVAS, and (F) fAAS. The shading represents hydrophobic interactions (green), hydrogen bond acceptors (blue); van der Waals contacts (grey residues); grey parabolas represent large accessible surface areas; thick lines around ligands illustrate accessible surface; and the size of the residue oval shapes represents the strength of the contact. Black curved lines highlight the side chain interactions for residues at position 1 (Met, Val and Ala, A—C, respectively).

A decrease in hydrophobic interactions is evident for fVAS, becoming more pronounced for fAAS relative to fMAS (Figure 2D, E, and F). Further, the overall docking values (Edock) correlated with van der Waals interaction (Egv) and hydrophobic potential (Egs) scores (Table S1-3), which demonstrated the highest variance between mutant peptides for the best poses, indicating that these interactions are likely the basis of substrate selectivity. The two-dimensional interaction diagrams (Figure 2D–F) shows a progressive decrease in accessible surface area for the side chain at position 1 of the peptides, which may explain the apparent decreased catalytic efficiency in hydrolysis of the formyl group of formyl valine bearing proteins, evidenced by their detection by mass spectrometry (Figure 1). It should be noted that the inability to remove the formyl group has been shown to compromise host viability^50,51^ and this would have potential implications if i-tRNA-AAC was used more widely to control essential genes within the cell, or to express a protein that relies on quick removal of the N-terminal residue for catalytic activity^52^.

### Overexpression of valyl-tRNA synthetase and methionyl-tRNA formyltransferase enhances translation initiation from the GUU start codon through increased i-tRNA-AAC charging and formylation

Previous work demonstrated that translation initiation from an i-tRNA-GAC mutant could be enhanced by the overexpression of translation-associated proteins^31^. Motivated by this and our observation of valine and formyl-valine initiated proteins, we investigated whether overexpression of ValRS and FMT could similarly improve translation initiation in the i-tRNA-AAC system. To test this, we engineered a set of plasmids to overexpress ValRS and FMT (Figure S9) and co-transformed them into strains harboring the core orthogonal i-tRNA-AAC translation initiation system (Core OTIS) (Figure 3A).

**Figure 3.**
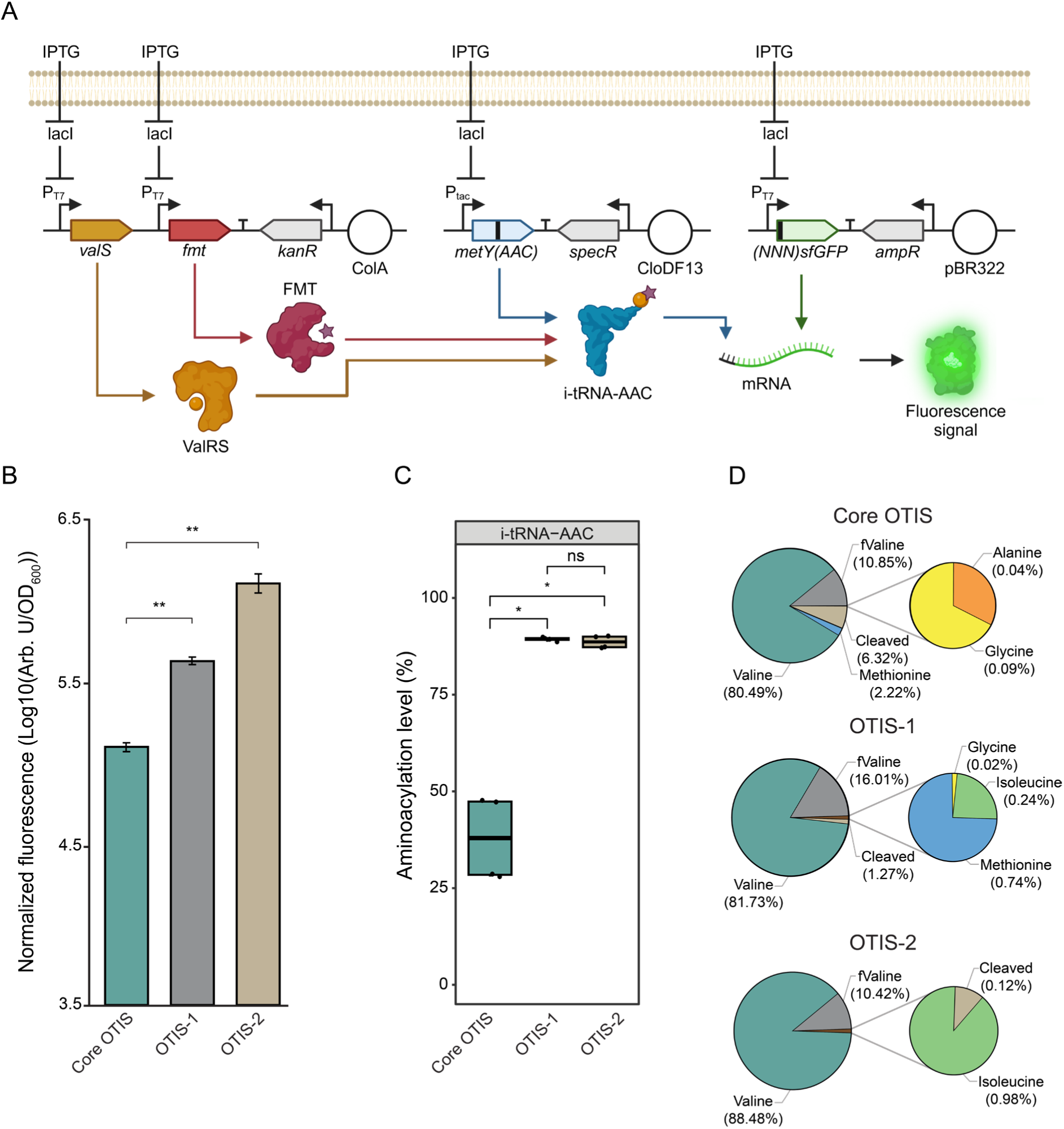
Overexpression of valyl-tRNA synthetase and methionyl-tRNA formyltransferase enhances translation initiation from the GUU start codon through increased i-tRNA-AAC charging and formylation. (A) Schematic overview of the i-tRNA-AAC mediated translation initiation system with ValRS and FMT co-overexpression. The mutant i-tRNA-AAC was expressed from an IPTG-inducible tac promoter on a pULTRA plasmid. Co-expressed is a start codon library variant encoding sfGFP reporter from the T7 promoter on a pET20B vector. ValRS and FMT proteins are also controlled by T7 promoters on a pCOLA vector. Created in BioRender. J, P. (2025) https://BioRender.com/a8iglz2 (B) Translation initiation from GUU start codon reporter is significantly increased OTIS-1 and OTIS-2 systems relative to Core OTIS. ANOVA p values, ** < 0.01. (C) Aminoacylation level of i-tRNA-AAC in Core OTIS, OTIS-1, and OTIS-2 systems. Wilcoxon signed-rank test p values, * < 0.05. (D) Proportion of amino acids used to initiate translation of (GUU)Nluc reporter from either the Core OTIS, OTIS-1, or OTIS-2 expression systems.

Overexpression of ValRS-alone alongside the Core OTIS (hereafter referred to as OTIS-1) significantly enhanced translation from sfGFP reporters bearing a GUU start codon by 3.3-fold (p value <0.01). Co-overexpression of both ValRS and FMT simultaneously (hereafter referred to as OTIS-2) had an additive effect, further increasing translation by an additional 3.0-fold relative to OTIS-1 (Figure 3B). In control strains lacking the Core OTIS, initiation from canonical AUG start codons were mildly reduced upon overexpression of ValRS or ValRS and FMT (Figure S10A), likely reflecting metabolic burden imposed by overexpressing additional heterologous proteins (Figure S10B). Collectively, these results suggest that lower aminoacylation efficiency of i-tRNA-AAC is a key bottleneck in Core OTIS performance, with inefficient subsequent formylation representing a secondary constraint.

To test whether improved aminoacylation of i-tRNA-AAC is a cause of the increased translation initiation observed in the OTIS-1 and OTIS-2 systems, we measured the tRNA charge levels in these strains using an adaptation of the mim-tRNAseq method. This method uses short-read next generation sequencing of reverse-transcribed periodate-treated extracted RNA to identify and differentiate aminoacylated and unaminoacylated tRNAs^53^. The mim-tRNAseq and associated computational toolkit have accurately measured tRNA abundance, aminoacylation levels, and modification status in yeast, fly, and human cells^54,55^. Using mim-tRNAseq, we found that OTIS-1 and OTIS-2 significantly increased the aminoacylation of i-tRNA-AAC by 1.6 – 2-fold (p value <0.01) (Figure 3C). These results confirm that inefficient aminoacylation is a primary constraint on Core OTIS performance and that this bottleneck can be partially relieved through ValRS overexpression.

Having increased i-tRNA-AAC-mediated translation initiation in OTIS-1 and OTIS-2, we next wanted to measure the impact this had on the proportions of N-terminally incorporated amino acids in proteins initiated with GUU start codons. We repeated the previous proteomic analysis on the Nluc reporter now expressed within OTIS-1 and OTIS-2 strains. Analysis of the OTIS variants revealed an inverse relationship between i-tRNA-AAC translation efficiency and the proportion of methionine-initiated proteins, which changed from 2.22% in the Core OTIS to undetectable levels in OTIS-2 (Figure 3D). The same was seen for cleaved peptide proportion (attributable to methionine removal by MAP) that reduced from 6.3% to 0.12% (Figure 3D). Similarly, small fractions of alanine (0.04%) and glycine (0.09%) initiated proteins were detected in the Core OTIS strain but were diminished or undetectable in OTIS-1 and OTIS-2 strains. Considering these findings alongside our detection of glycine-initiated proteins in the (AUG)Nluc control (Figure S4), points to their incorporation from native i-tRNA-fMet initiation. These data suggest that improved i-tRNA-AAC initiation efficiency, resulting from a higher fraction charged with formylated valine, reduced background translation from GUU start codons by the native initiator tRNA. By contrast, proteins initiated with isoleucine were observed only in the OTIS-1 (0.24%) and OTIS-2 (1%) strains (Figure 3D), suggesting they may have been incorporated by mischarged i-tRNA-AAC.

### Overexpression of valyl-tRNA synthetase (ValRS) suppresses aminoacylation of specific endogenous tRNAs while down-regulating acp^3^U47 modification on valine isoacceptors

Having characterized and improved the orthogonal i-tRNA-AAC-mediated translation system, we next wanted to determine the impact of deploying OTIS-1 and OTIS-2 on host tRNA transcript abundance, aminoacylation levels, and base-modifications using mim-tRNAseq. These measurements showed OTIS-1 did not significantly change tRNA transcript abundance, while OTIS-2 resulted in 4.5-5-fold (p value < 0.01) increases in tRNA-Val-GAC-1 and tRNA-Val-GAC-2 transcripts (Figure S11). Aminoacylation levels for the majority (73%) of the endogenous tRNAs did not change between wild-type and the OTIS variants (Figure S12). Of those that did change, tRNA-Gln-CUG-1 and tRNA-Leu-CAG-1 aminoacylation levels were slightly increased in Core OTIS and further increased by OTIS-1 and OTIS-2 (Figure 4A). By contrast, the aminoacylation levels of tRNA-Ala, tRNA-Ser, and tRNA-Thr isoacceptors significantly decreased in OTIS-1 and OTIS-2.

**Figure 4.**
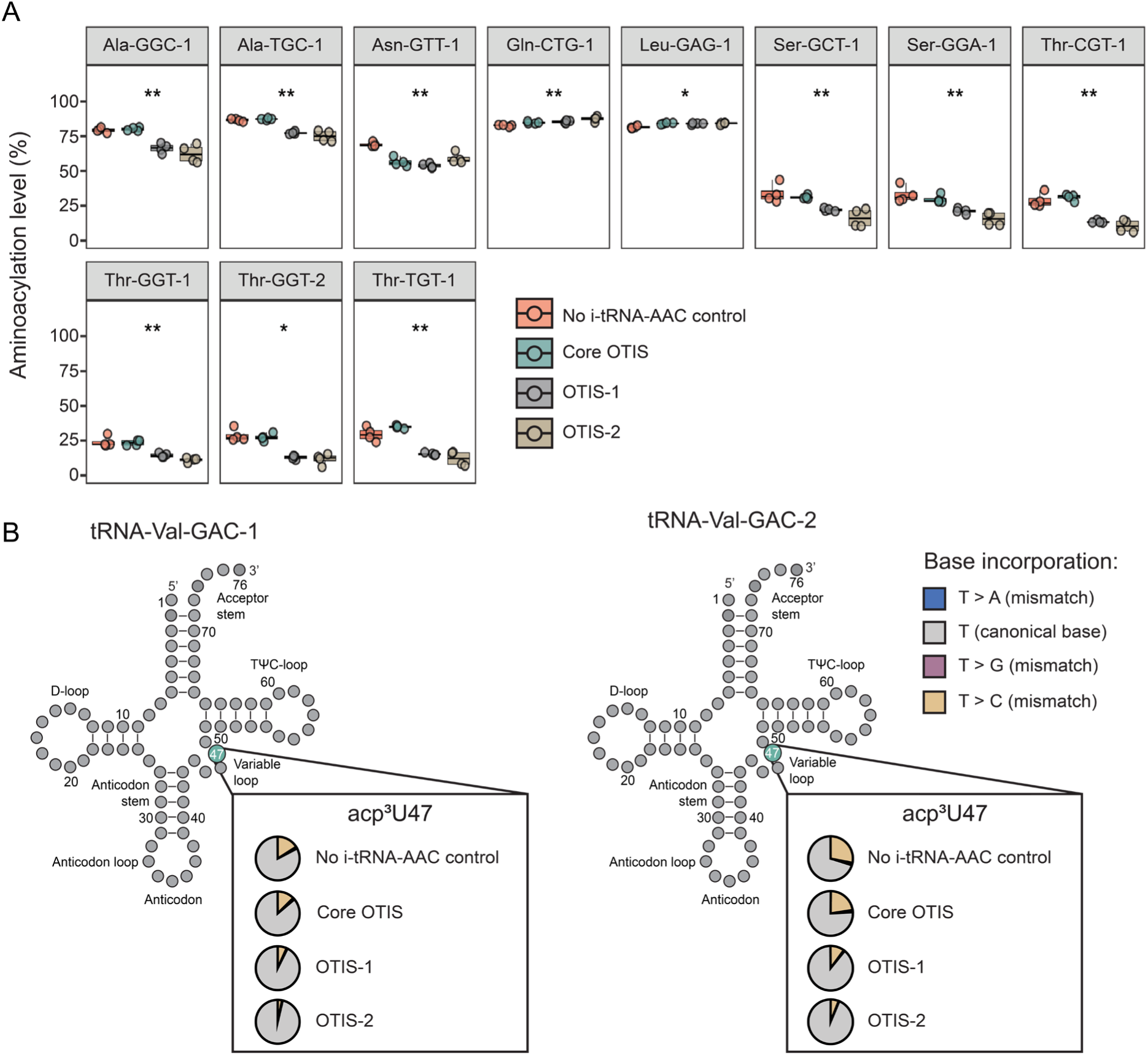
**Overexpression of valyl-tRNA synthetase (ValRS) suppresses aminoacylation of specific endogenous tRNAs while down-regulating acp^3^U47 modification on valine isoacceptors**. (A) Mim-tRNAseq measurements of aminoacylation level of affected endogenous tRNAs upon deployment of Core OTIS, OTIS-1, or OTIS-2 systems. (B) Pie charts represent the average reverse transcriptase (RT) base incorporation frequency at position 47 in Val-GAC tRNA isodecoders (tRNA-Val-GAC-1 and tRNA-Val-GAC-2). Both aminoacylation level and RT base incorporation frequency was measured from strains harboring the Core OTIS, OTIS-1, or OTIS-2 systems and compared to no i-tRNA-AAC control strains. tRNAs named according to GtRNAdb convention, omitting the Gene locus ID as these are all identical gene sequences that are indistinguishable using our sequencing method ^58^. Kruskal Wallis H-Test p values for aminoacylation levels, *** < 0.001, ** < 0.01, and * < 0.05.

Base modifications of tRNAs are important for their structure, stability, fidelity, and rate of translation^56^. We reasoned that while mim-tRNAseq was originally designed and validated on eukaryotic tRNAs^54,55^, the modification-induced reverse transcriptase (RT) misincorporations used as signatures of modification sites should still be robustly generated by prokaryotic base modifications. Using mim-tRNAseq we measured stable reverse-transcription misincorporation rates at predicted modification sites (annotated in the MODOMICS database) across the endogenous tRNAs (Figure S13). Comparing the rate of misincorporation at known tRNA modification sites, we observed one major deviation from this pattern at position U47 of both tRNA-Val-GAC isodecoders (Figure 4B). We consistently observed TGIRT reverse transcriptase misincorporation of U47C at this site progressively decreased from Core OTIS to OTIS-2. In *E. coli* tRNA-Val-GAC isodecoders, position 47 contains an 3-(3-amino-3-carboxypropyl)uridine (acp^3^U) modification important for structural stability^57^. Although several other *E. coli* tRNAs (e.g., tRNA-Arg-ACG, tRNA-Ile-GAU, tRNA-Ile-CAU, tRNA-Met-CAU, tRNA-Lys-UUU, and tRNA-Phe-GAA) showed similar U47C misincorporation by TGIRT, the magnitude of the misincorporation changes did not show a similar trend (Figure S13).

### Overexpression of valyl-tRNA synthetase and methionyl-tRNA formyltransferase induces global proteomic perturbations

To understand global effects on the *E. coli* proteome from OTIS-1 and OTIS-2, we carried out proteomic profiling using a data-independent acquisition (DIA) mass spectrometry workflow (Figure S14). The Core OTIS was well tolerated, with no significant proteomic perturbations observed (Figure 5A), in contrast to OTIS-1 or OTIS-2 that caused significant proteome remodeling. Deployment of OTIS-1 resulted in 19 up-regulated and 28 down-regulated proteins (Figure 5B), while OTIS-2 induced more significant changes, with 76 proteins up-regulated and 216 down-regulated (Figure 5C). Gene ontology enrichment analysis revealed that in OTIS-2, proteins associated with the tricarboxylic acid (TCA) cycle and fatty acid beta-oxidation were broadly downregulated, whereas components of the fatty acid biosynthesis pathway were upregulated (Figure 5D). Despite these changes, canonical stress responses such as the stringent response^59^ were not activated, suggesting that the host maintains significant metabolic flexibility.

**Figure 5.**
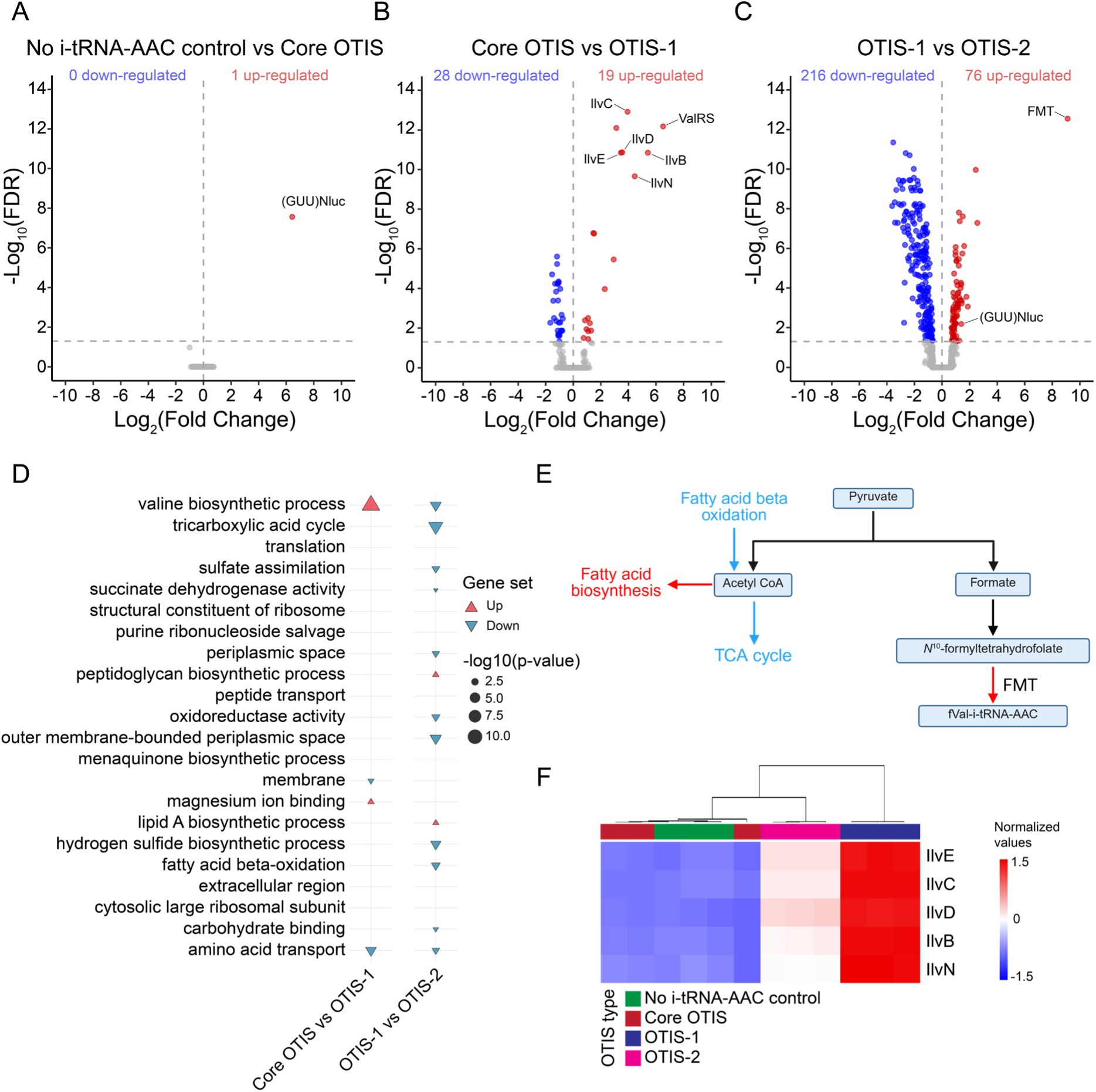
Overexpression of valyl-tRNA synthetase (ValRS) and methionyl-tRNA formyltransferase induces global proteomic perturbations. Volcano plots of differentially expressed proteins between (A) No i-tRNA-AAC control vs Core OTIS, (B) Core OTIS vs OTIS-1, and (C) OTIS-1 vs OTIS-2. Proteins with significant increases in abundance are shown in green (FDR < 0.05, Log2 fold-change > 0) and proteins with significant decreases in abundance are shown in blue (FDR < 0.05, Log2 fold-change < 0). (D) Gene set enrichment analysis of differentially expressed proteins in Core OTIS, OTIS-1, and OTIS-2 systems. Enriched pathways are grouped by functional annotation with up and down regulated functional annotations indicated by triangle orientation -log10(p-value) represented by the size of the triangles. (E) Proposed model linking increased demand of i-tRNA-AAC formylation to host metabolism. Pathways colored in red are up regulated and pathways colored in blue are down regulated. (F) Heatmap of differentially expressed proteins involved in the valine biosynthesis pathway. Samples were hierarchically clustered based on their expression profiles, and relative protein fold changes are shown as normalized values ranging from -1.5 to 1.5, represented by a blue to red color scale.

Consistent with this flexibility, the expression levels of IlvB, IlvC, IlvD, IlvE and IlvN proteins associated with the isoleucine/valine biosynthetic pathway were robustly upregulated in both OTIS-1 and OTIS-2, possibly as an adaptive response to increased demand for valine being charged to i-tRNA-AAC (Figure 5F). Despite the magnitude of valine pathway upregulation being attenuated in the OTIS-2 condition relative to OTIS-1, these proteins remained upregulated compared to the Core-OTIS, reflecting how increasing translational burden may limit the hosts’ native adaptive potential. Taken together, these findings highlight the capacity of the host to dynamically rewire its proteome and metabolism to support orthogonal translation systems. At the same time, our results also reveal that resource constraints imposed by extensive protein overexpression can limit otherwise adaptive metabolic responses, reinforcing the importance of balancing system output with host burden during the design and deployment of these engineered systems^34,60,61^.

## Discussion

In this study, to deeply understand the interaction between an engineered i-tRNA and its host, we developed a multi-omic analytical pipeline. Using this pipeline, we show the variety of amino acids that could be incorporated at the N-terminus by both native and engineered i-tRNAs along with their formylated variants (Figure 1). Additionally, we use *in silico* modeling to investigate a plausible mechanism for the observed preservation of the formyl group on N-terminally incorporated valine (Figure 2). Our measurements of aminoacylation efficiency (Figure 3) enabled us to relieve a bottleneck limiting i-tRNA-AAC-mediated protein translation by expression of accompanying ValRS and FMT, proving the utility and mechanistic insight of this approach. Further, this translation efficiency improvement also appeared to reduce background translation from the GUU start codon by native i-tRNA, underscoring the importance of efficient tRNA charging in maintaining orthogonality^27,62,63^.

### Charging and formylation are the dominant bottlenecks in initiation

Multi-omics measurements converged on two rate-limiting steps for i-tRNA-AAC-mediated initiation: aminoacylation of i-tRNA-AAC by ValRS and formylation of the valyl-charged initiator by FMT. We chose to overexpress ValRS to overcome inefficient aminoacylation based on three factors: (1) previous work showing chimeric tRNAs possessing mixed aaRS determinants are inefficiently aminoacylated^62,63^; (2) reporter proteins initiated by i-tRNA-AAC predominantly possessed N-terminal valine (Figure 1E); and (3) the A35 and C36 nucleotides present in the AAC anticodon are strong identity determinants for ValRS recognition^33^. Using fluorescence assays and mim-tRNAseq, we clearly showed an increase in translation initiation from i-tRNA-AAC when overexpressing ValRS to increase the proportion of charged i-tRNA, demonstrating the utility of the approach^62^.

A second avenue for improving the aminoacylation efficiency of the mutant i-tRNA-AAC may lie in enhancing its intrinsic recognition by endogenous ValRS. However, pursuing this strategy is challenging, as many of the sequence and structural features that influence ValRS mediated aminoacylation overlap with the elements required for proper initiator tRNA functioning. For example, it has previously been shown that increasing the rigidity of the anticodon stem of Val-tRNA by mutating the U29:A41 pair to C29:G41 greatly diminishes aminoacylation efficiency^64^. Therefore, the highly conserved 3GC motif present in our chimeric i-tRNA-AAC, essential for its role in initiation as it licenses transitioning from the 30S initiation complex to the elongation competent 70S complex^24^, may simultaneously be hindering its aminoacylation. Although previous work has shown that the middle G30:C40 pair of the 3GC motif was critical and that initiation can tolerate limited mutations in the flanking GC pairs, these flanking pairs remain necessary to achieve optimal initiation efficiency^24^. Therefore, while modifying this anticodon stem feature could in principle improve ValRS recognition, such changes are constrained by the structural requirements necessary for the tRNA to function as a competent initiator. Overcoming these constraints will likely require approaches beyond rational design that leverage high-throughput strategies, such as Chi-T^65^, to screen large numbers of tRNA variants. Doing so may uncover i-tRNA sequences not readily predictable from existing structural tRNA rules and rapidly accelerate the development of OTSs. While aminoacylation represents a primary bottleneck in i-tRNA-AAC mediated initiation, a second rate-limiting step arises after charging, namely the efficient formylation of the valine bearing i-tRNA.

Although formylation of aminoacylated i-tRNAs is not strictly required for translation initiation, it enhances initiation efficiency by increasing the affinity of the charged i-tRNA for initiation factor 2 and promotes its recruitment to the 30S ribosomal subunit^50,63^. We observed this effect in OTIS-2, which displayed a 3-fold increase in translation efficiency relative to OTIS-1 (Figure 3). However, an *in vitro* study showed that FMT exhibits a ∼60-fold lower catalytic efficiency V_max_ when acting on valine-charged i-tRNAs compared to their canonical methionine counterparts^66^. Inefficient formylation of valine-bearing i-tRNAs was therefore likely a key contributor to the reduced initiation efficiency. However, previous work by Shetty *et al*.^24^ showed that overexpression of i-tRNA-fMet in a formylase-deficient mutant could partially compensate for the absence of formylation, suggesting higher levels of i-tRNA can offset, but not fully overcome poor formylation. This likely explains the improvement seen in OTIS-1, with further gains achieved after overexpression of FMT increased the extent of i-tRNA-AAC formylation. Supporting this notion, the proportion of aminoacylated i-tRNA-AAC did not change between OTIS-1 and OTIS-2 (Figure 3C), whereas translation initiation efficiency increased markedly (Figure 3B).

### Crosstalk between ValRS and i-tRNA-AAC with the host tRNAome

We show that the overexpression of the translation associated enzymes, ValRS and FMT, not only improved orthogonal translation initiation efficiency but also enhanced system fidelity by suppressing the competing native initiation system. This was evidenced by the reduction in methionine incorporation at GUU start codons from 2.22% to undetectable levels (Figure 3D). Despite these functional gains, our multi-omic analysis reveals that increasing the expression of these enzymes produces modest effects on the host tRNAome. Global tRNA transcript abundance, aminoacylation levels, and modification profiles remained largely unchanged, with only a small subset of tRNAs exhibiting changes (Figure 4). Unsurprisingly, these findings suggest that although reengineering of OTISs can improve orthogonality and performance, it can also simultaneously introduce crosstalk within the tRNA landscape.

A notable example of this crosstalk emerged from a potential mischarging event, shown by the unexpected appearance of isoleucine-initiated proteins in OTIS-1 (0.24%) and OTIS-2 (1%) (Figure 3D). This contrasts with the suppression of the use of methionine at GUU codons for initiation and instead increases as OTIS output increases. Previous *in vitro* studies have demonstrated that ValRS is capable of stably mischarging valine isoacceptors with isoleucine as the amino acid is readily accommodated into the activation site while being sterically excluded from the separate editing domain^67^. In a native cellular context, this mischarging of valine isoacceptors are considered infrequent owing to the comparatively low abundance of isoleucine versus valine and the higher affinity of ValRS for its cognate amino acid^67,68^. However, elevating the pool of ValRS in OTIS-1 and OTIS-2 could relax this substrate competition and enable the stable mischarging of the engineered i-tRNA-AAC with isoleucine. Subsequent initiation at GUU start codons with this i-tRNA-AAC-Ile would then result in isoleucine incorporation at the N-terminus in our reporter proteins (Figure 3D).

A second class of tRNA perturbation identified by mim-tRNAseq analysis involved reduced aminoacylation levels of endogenous alanine-, serine-, and threonine-bearing tRNAs in OTIS-1 and OTIS-2 (Figure 4A). As with isoleucine misincorporation, ValRS overexpression may contribute to this effect. However, unlike isoleucine, the amino acids alanine, serine, and threonine are efficiently accommodated into the editing site of ValRS, preventing their misincorporation into proteins^67^. We speculate that upon ValRS overexpression, the frequency of this non-cognate mischarging may increase, leading to the transient sequestration of alanine, serine, and threonine by ValRS. Further, this effect may be exacerbated by the post-transfer editing dominant nature of ValRS, as the release of non-cognate substrates from the editing domain is dependent on the presence of cognate tRNA-Val^64,69–71^. This prolonged residency may diminish the free pools of these amino acids available to their cognate synthetases, producing the observed reductions in aminoacylation levels.

The measured tRNA epitranscriptome remained broadly stable upon deployment of Core-OTIS, with only a single notable deviation emerging in OTIS-1 and OTIS-2 (Figure 4B). Specifically, we observed a progressive reduction in the U47C misincorporation signature on both tRNA-Val-GAC isodecoders, corresponding to the known site of the acp^3^U modification involved in structural stability^57^. This suggests that while the overexpression of ValRS and FMT improved i-tRNA-AAC mediated translation, it selectively influenced the modification state of valine elongator tRNAs. Although several other tRNAs also displayed U47C misincorporation at this position (Figure S13), none exhibited a comparable directional change across OTIS variants, indicating that this effect was selective rather than global. This specificity toward valine tRNAs was consistent with their cognate interaction with the overexpressed ValRS pool and the unique demands placed on valine tRNA processing. This result highlights that despite the tRNA epitranscriptome being broadly stable, orthogonal translation systems can impose highly localized pressures on specific tRNA species, potentially impacting the functional behavior of the affected tRNAs in ways that may influence both engineered and native translation processes.

### Proteome-wide remodeling reflects resource allocation and metabolic demand

Overexpression of i-tRNA-AAC, ValRS, and FMT in OTIS-2 did not have large-scale off-target effects on cellular tRNA transcript levels (Figure S11), aminoacylation (Figure S12), or epitranscriptome (Figure 4). However, larger effects were seen at the proteome level (Figure 5) with broad remodeling in energy metabolism and biosynthetic pathways. In total, as many as 292 proteins were found to be differentially expressed, representing ∼14% of the quantified proteome, similar in magnitude to other metabolic shifts such as mild acid stress^72^. One possible mechanism underlying this metabolic reprogramming could be an increased flux through pyruvate catabolism, which generates formate to support the increased demand for i-tRNA-AAC formylation (Figure 5E). The concurrent production of acetyl-CoA as a byproduct may contribute to the observed upregulation of lipid biosynthesis, potentially serving as a mechanism for balancing metabolic supply. Alternatively, this shift may reflect a broader anabolic remodeling to accommodate the increased proteomic load associated with the overexpression of proteins and to preserve cellular morphology and structural integrity.

These measurements underscore the careful balance that must be struck between increasing the efficiency of engineered translation systems and inadvertently introducing physiological burdens on the host^22,34,73^. This work is a step towards having the analytical tools in place to measure and respond to these perturbations.

### Design principles and opportunities for next-generation orthogonal translation initiation

In synthetic biology, the capacity to precisely control translation initiation could facilitate the advancement of genetic code expansion efforts^74–76^ and engineering of increasingly sophisticated orthogonal synthetic circuits^5,40^. In this space, i-tRNA mutants capable of initiating translation from non-canonical start codons have emerged as a promising tool^19–22^. However, the utility of such systems hinges on their ability to operate with minimal interference from endogenous translational machinery and other co-expressed synthetic genetic circuitry. For this reason, deeper mechanistic characterization of orthogonal systems and their effects on the host are essential, in particular when creating codon-reduced organisms with less redundancy in their genetic code^77–79^.

Future work seeking to leverage orthogonal translation systems could implement the systematic identification of these perturbations and employ data-driven optimization to balance system performance while minimizing undesirable physiological trade-offs^77,80^. Through carefully tuning and optimizing these molecular interactions, we can not only mitigate host burden but also significantly expand these tools toward broader applications^81^. For example, the i-tRNA-AAC could be the foundation for a future multiplexed translation initiation platform, particularly when integrated with strategies that allow independent regulation of multiple i-tRNAs^82^ and the tuning of downstream protein expression timing, abundance, and turnover through the N-degron pathway^8,19,82^.

Overall, this study offers unique insights into the biochemical and cellular implications of engineering OTISs. By quantifying system orthogonality, identifying rate limiting steps and mapping host responses, we provide a framework for the rational design of i-tRNA based OTSs and host compatibility assessment for future designs.

### Limitations of the study

Although our study demonstrated the successful deployment and mechanistic characterization of an orthogonal translation initiation system based on i-tRNA-AAC, several limitations should be noted. All measurements were performed in the BL21(DE3)pLysS strain, which may restrict the generalizability of our findings to other strains with different tRNA pools, metabolic profiles, or stress-response networks. Furthermore, while our structural docking analysis provides a plausible explanation for the reduced efficiency of formyl-valine deformylation, this hypothesis requires validation through *in vitro* or *in vivo* kinetic studies. We also focused exclusively on overexpressing ValRS; although other aminoacyl-tRNA synthetases could theoretically aminoacylate i-tRNA-AAC, mass spectrometry data suggest this would likely offer limited benefit given valine’s dominance as the N-terminal residue. Additionally, we cannot definitively identify which tRNA species delivered the amino acids observed at the N-termini of reporter proteins in our mass spectrometry analysis, and further investigation is needed to resolve the tRNA identity for each initiation event. Finally, our implementation of the mim-tRNAseq method using TGIRT and its associated analysis pipeline captures only a subset of tRNA modifications, constraining the completeness of our epitranscriptomic profiling.

## Materials and Methods

### Bacterial Strains

All plasmids within this study were expressed in BL21(DE3) (ATCC #BAA-1025) or BL21(DE3)pLysS strains (Promega, Cat. No. L1195). All strains were grown in lysogeny broth Miller (10 g/L tryptone, 10 g/L NaCl, 5 g/L yeast extract and Milli-Q® water) (LB^M^) or on LB^M^ agar (10 g/L tryptone, 10 g/L NaCl, 5 g/L yeast extract, 15 g/L agar and Milli-Q® water), supplemented with appropriate antibiotics. Chloramphenicol (25 µg/mL) was used in the media for BL21(DE3)pLysS growth, spectinomycin (50 µg/mL) for strains containing a pULTRA plasmid, carbenicillin (100 µg/mL) for strains containing pET20b plasmid, and kanamycin (50 µg/mL) for strains containing the pCOLA plasmid. In strains harboring four plasmids, four antibiotics were used but at half-standard concentration. All bacterial cultures within this study were inoculated into 2 mL of LB^M^ with appropriate antibiotics in 15 mL conical tubes and grown overnight (16 hours) in an Infors MT Multitron pro-orbital shaker-incubator at 250 RPM rotating orbitally at 25 mm diameter at 37°C, unless stated otherwise. These stationary phase overnight liquid cultures were back diluted to 0.1 OD_600_ and grown in 15 mL conical tubes in an Infors MT Multitron pro-orbital shaker-incubator at 250 RPM rotating orbitally at 25 mm diameter at 37°C, unless stated otherwise. Bacterial growth parameters have been reported to the best of our knowledge conforming with the MIEO v0.1.0 standard^83^.

### Plasmid Construction

To create the pULTRA-*tac*-*metY(AAC)* plasmid we designed a mutated version of *metY* from MG1655 (GenBank: U00096.3) where the anticodon was changed from CAT to AAC (Table S5). The pULTRA-CNF vector, which was a gift from Peter Schultz (Addgene plasmid # 48215)^84^, was linearized by inverse PCR (Table S6) and ligated with synthetic DNA containing the *metY* gene or the mutated *metY*(AAC) (Table S5) genes using Gibson assembly (NEBuilder HiFi DNA Assembly kit (NEB#E2621)). The pULTRA-*tac*-*Empty* control plasmid was constructed by re-ligating the linearized pULTRA-*tac*-*metY(AAC)*, removing the *metY*(AAC) gene in the process. The pET20b-*T7-(NNN)sfGFP* reporter plasmids were from previous work by Hecht *et al.* ^37^. Using similar methods to construct the pULTRA vector, the pET20b-*T7-(NNN)Nluc6His* was built using the template pET20b-*T7-(AUG)sfGFP* and synthetic DNA containing the *nluc* gene variants (Tables S5 and S6). The pCOLA plasmid variants were generated using the pCOLA-*T7-WspR* template plasmid which was a gift from Ming Hammond (Addgene plasmid # 79162) ^85^. The WspR gene was removed during vector linearization through inverse PCR (Table S6). *Fmt* and *ValRS* genes were amplified from the genome of *E. coli* BL21(DE3)pLysS (Table S6) and were then cloned into the linearized pCOLA vectors through Gibson assembly. To generate the pCOLA-*T7*-*ValRS/fmt* plasmid, this process was repeated using the newly constructed pCOLA*-T7*-*ValRS* plasmid as a template (Table S6). The pCOLA-*T7*-*Empty* control plasmid was generated through the re-ligation of the linearized pCOLA vector. The final constructs were verified through whole plasmid sequencing (Plasmidsaurus).

### Bacterial Fitness Analysis

Conducted as previously^22,23^ with modification. Briefly, BL21(DE3)pLysS strains containing either plasmid pULTRA-*tac*-*metY(AAC)* or pULTRA-*tac*-*Empty*, were grown overnight at 37°C with shaking in 2 mL of LB^M^ with the addition of spectinomycin and chloramphenicol. Overnight cultures of these strains were passaged and diluted 1:100 into 2 mL of fresh LB^M^ and the appropriate antibiotics and grown at 37°C with shaking until cultures were at 0.6 OD_600_. Following this, cells were back diluted to 0.1 OD_600_ with a final volume of 200 µL of LB^M^ and appropriate antibiotics in a flat bottom 96 well plate (Sigma, cat#P8116) and sealed with a gas-permeable transparent seal (Sigma cat# Z380059). Within this plate, two different conditions were set up: an induced condition containing LB^M^ with the appropriate antibiotics and 1 mM IPTG and a control condition which only contained LB^M^ and the appropriate antibiotics. Cells were grown for 7 hours at 37°C in a SPECTROStar Nano plate reader, with absorbance at OD_600_ readings occurring every 5 minutes. Growth rate (μr) and maximum cell optical density (maxOD_600_) were analyzed using the GrowthCurver R package^86^.

### Fluorescence Measurements

*E. coli* BL21(DE3)pLysS strains harboring the relevant plasmids were grown overnight at 37 °C in 2 mL of LB^M^ in the presence of carbenicillin, spectinomycin, chloramphenicol, or kanamycin, as required, in 96 deep well plates (Sigma, cat#P8116). Bacterial cultures were then diluted 1:100 in 400 µL of fresh LB^M^ and appropriate antibiotics and grown for 2 hours at 37 °C with shaking. The cultures were then induced with 1 mM IPTG, and growth for a further 4 hours at 37 °C, or until cells reached 0.6 OD_600_, in preparation for bulk fluorescence measurements and flow cytometry. Following induction, 400 µL of culture was transferred into a clear 96-well plate (Sigma, cat#CLS3610) and centrifuged in a swinging bucket rotor at 2,240 x *g* for 12 minutes at 20°C. Following this, the supernatant was removed, and the pelleted cells were resuspended in 200 µL of phosphate-buffered saline (PBS) and transferred into a black clear bottom 96-well plates (Sigma, cat#CL3603). Resuspended cells were stored at 4°C overnight to allow for sfGFP maturation^37^. Absorbance at OD_600_ and bulk fluorescence for sfGFP expression was measured using the PHERAstar microplate reader (BMG Labtech) (excitation 488 nm, emission 530 nm). Results were recorded with a gain setting of 200. Normalized (Arb. fluorescence units/OD_600_) results were analyzed using the DEseq2 R package^38^. Parameters were set to find fluorescence changes of 4-fold or more, with a statistical significance of p-value <0.01.

### Flow Cytometry

Bacterial cultures were prepared as above for fluorescence measurements. Following this, cells were diluted 100-fold in PBS to reach the final cell concentration of approximately 10^6^ cfu/mL and measured on a CytoFLEX S (Beckman Coulter) using a FITC fluorescence channel with a 488 nm excitation laser and 525/40 nm emission band-pass filter with side scatter triggered events threshold set to 10,000 events. Data processing and analysis were performed using CytExpert (Version 2.3.0.84) (Beckman Coulter) and the DEseq2 R package^38^. Parameters were set to find fluorescence changes of 4-fold or more, with a statical significance of *p<0.01*.

### Expression and Purification of Nluc

Biological triplicates of single colony BL21(DE3)pLysS cells harboring the pULTRA-*tac*-*Empty* or pULTRA-*tac-metY(AAC)* plasmids, the pET20b-*T7-(AUG)Nluc6HIS* or pET20b-*T7-(GUU)Nluc6HIS*, or one of the pCOLA*-T7-Empty*, pCOLA-*T7-ValRS*, or pCOLA*-T7-ValRS/fmt* plasmids were used to inoculate 5 mL of LB^M^ supplemented with spectinomycin, chloramphenicol, carbenicillin, or kanamycin as needed and incubated at 37°C overnight with shaking. The protocol from Hecht *et al.* ^37^ was used for expression and purification of Nluc with the following modifications. The overnight cultures were diluted 1:100 in 30 mL of LB^M^ with appropriate antibiotics and incubated at 37°C for 2 hours, followed by induction with 1 mM of IPTG and a final outgrowth step for an additional 4 hours to allow for sufficient protein expression. The cell cultures were pelleted at 3,500 x*g* for 10 min. The supernatant was discarded, and the cell pellets were chemically lysed using 400 µL lysis buffer (CelLytic B cell lysis reagent (Sigma-Aldrich #B7310), 0.1 mg/mL lysozyme, 1x protease inhibitor (Roche #04693132001), and 100 units of benzonase (Sigma-Aldrich #E1014)). The resuspended cells were incubated at room temperature for 20 min to allow for sufficient lysis. The soluble protein fraction was then collected by centrifuging the cell lysate at 16,000 x*g* for 10 minutes. The expressed C-terminal HIS-tagged Nluc was Ni-NTA purified using HisPur^TM^ Ni-NTA Resin (ThermoFisher Scientific, #88221) as per the HisPur^TM^ Ni-NTA Resin protocol. Equilibration, wash, and elution buffers were made using 300 mM PBS with 20 mM, 60 mM, and 500 mM imidazole, respectively.

### Data Dependent Acquisition Mass Spectrometry

Purified Nluc samples were reduced through the addition of dithiothreitol (DTT) to a final concentration of 10 mM and incubated on a heat block at 60°C for 30 minutes. Samples were alkylated using iodoacetamide at a final concentration of 30 mM and incubated at room temperature in the dark for 1 hour. Purified Nluc was precipitated out of solution using 1 mL of ice-cold acetone incubated overnight at -20°C and pelleted via centrifugation at 16,000 x*g* for 10 min at 4°C. The supernatant was decanted, and the protein pellet was left to dry for 15 minutes at room temperature in a chemical fume hood. The dried protein pellet was resuspended in 8 M Urea, 50 mM Tris-HCl buffer (pH 8.0) and diluted 5-fold by addition of 50 mM Tris-HCl buffer (pH 8.0). Protein samples were trypsin digested using a 1:50 (w/w) ratio and incubated at 37°C overnight. Peptides were then acidified with 1% (v/v) formic acid, followed by C18 stage tip purification (Merck, #66883-U). The peptides were then dried in a SpeedVac vacuum concentrator and reconstituted to a final concentration of 1 μg/μL in solvent A (2% (v/v) acetonitrile / 0.1% (v/v) formic acid).

Samples were separated using a Dionex nanoLC system and analyzed on a high-resolution Q-Exactive HF-X mass spectrometer (Thermo Fisher Scientific) using a DDA method with an inclusion list (File S1). Reconstituted peptides were injected onto a C18 reverse phase column and eluted over a 70-minute linear gradient by increasing concentrations of 2-35% solvent B (99.9% (v/v) acetonitrile, 0.1% formic acid) relative to solvent A. An initial survey scan was conducted at 120,000 resolution with a scan range of 350 to 1,650 *m/z*, AGC target set to 3 x 10^6^, and a maximum injection time of 50 ms. The inclusion list used for N-terminal peptides (File S1) was generated using Skyline^87^ (Version 23.1.0.268) with general dd settings set to if idle, pick others. Precursors ions were subjected to high collision dissociation fragmentation using a normalized collision energy of 27. Resulting product ions were analyzed in an MS/MS scan set at 30, 000 resolution, AGC target of 1 x 10^5^, maximum injection time of 100 ms, and a 1.4 m/z isolation window. Analysis of precursor and product ion spectra was performed using MSFragger (Version 3.8) ^88^ and Skyline (Version 23.1.0.268)^87^.

### Data-Independent Acquisition Mass spectrometry

The whole proteome from *E. coli* BL21(DE3)pLysS cells harboring the pULTRA-*tac*-*Empty* or pULTRA-*tac-metY(AAC)* plasmids, the pET20b-*T7-(GUU)Nluc6HIS*, and one of the pCOLA-*T7-Empty*, pCOLA-*T7-ValRS*, or pCOLA-*T7-ValS/fmt* plasmids were prepared as described above without carrying out the Ni-NTA purification step. Samples were then separated using a Dionex nanoLC system and analyzed on a high-resolution Q-Exactive HF-X mass spectrometer (Thermo Fisher Scientific) using a DIA method. Reconstituted peptides were injected onto a C18 reverse phase column and eluted over a 150-minute linear gradient by increasing concentration of 2-35% solvent B (99.9% (v/v) acetonitrile, 0.1% formic acid) relative to solvent A (2% (v/v) acetonitrile, 0.1% formic acid). An initial precursor scan was conducted at 120,000 resolution with a scan range of 350 to 1,650 *m/z*, AGC target set to 3 x 10^6^, and a maximum injection time of 50ms. The subsequent scans were carried out using a 20-variable window scheme covering the precursor mass range of 350 to 1650 *m/z* at 30,000 resolution with an AGC target set to 2 x 10^5^ and normalized collision energy of 27.5.

Raw files were then analyzed against a FASTA file of *Escherichia coli* BL21(DE3) UniProtKB database using DIA-NN (Version 1.9.1) ^89^ to detect and quantitate peptides. The precursor *m/z* range was from 300 to 1800 *m/z,* and fragment ion range set from 200 to 1800 *m/z* with a peptide length range of 7 to 30. N-term M excision, C carbamidomethylation, and methionine oxidation were included as fixed and variable modifications, and a 1% precursor FDR was applied. All other DIA-NN parameters were set to default.

### Data filtering and normalization

The datasets were filtered using peptide spectral matches (PSMs) level data, retaining only entries with more than 2 PSMs. PSMs were considered distinct if they had different sequences or if they had the same sequence but differed in their post-translational modifications. PSMs were then summed to peptide level, and peptides with more than 50 percent of the samples with missing values or values below the first percentile of intensity values in each group for more than 50 percent of the experimental groups were discarded from the analysis. Peptides were rolled up to protein-level data using the IQ R package (https://github.com/tvpham/iq) without normalization. Zero values were replaced with NA denoting these are missing values in the resulting protein intensity matrix. Proteins with more than 50 percent of the samples with missing values or values below the first percentile of intensity values in each group for more than 50 percent of the experimental groups were discarded from the analysis. The data matrix was then log (base 2) transformed and between-sample normalization was performed using the cyclic LOESS method from the ‘limma’ R package^90^. To remove potential batch effects from biological data contained in the normalized data matrix, we used (RUVIII-C) R package^91,92^ to remove unwanted variation. The method relies on having a set of endogenous negative control proteins, which are proteins with little or no changes in protein abundances between different samples. For this study, 100 empirical negative control proteins (q-value < 0.05) were identified from an initial ANOVA test comparing across the biological replicate groups. Four unwanted components were selected for removal. At each stage of the data normalization, the samples were checked for batch effects using principal component analysis (PCA) plot and relative log-expression (RLE) plots and the distribution of the Pearson correlation of every pair of biological replicate samples were calculated.

### Differential abundance and pathways enrichment analysis

Differential abundance analysis of proteins was performed using the adjusted abundance matrix, to compare each group using the ‘limma’ R package ^90^. The linear model was fitted using the ‘lmFit’ function and the p-values calculated using the empirical Bayes method ‘eBayes’ function with trended and robust analysis enabled. The false discovery rate correction was applied to the moderated p-values by calculating the q-values ^93^. The treat function was used to find proteins with high confidence log fold-change above the threshold of 1.5. Significant differentially expressed proteins were defined as those with q-values less than 0.05. GO annotations were downloaded from the UniProt database (https://www.uniprot.org/proteomes/UP000002032). GO pathways enrichment was performed using the ClusterProfiler R^94^ package with all proteins that passed quality checking included as background. Pathways that had an adjusted p-value of < 0.05 were included for downstream analysis. The Gene Ontology (GO) enrichment results were further processed for visualization as a dotplot matrix. The normalized enrichment score was computed as the log2 ratio of the background gene count to the gene count in the enriched set.

### Protein:protein Interaction Stability and Docking Analyses

Peptide binding stability and optimal docking of substrate tripeptides were calculated in Molsoft ICM version 3.9-2a^95^ using the pdb-redo^96^ refined structure of *E. coli* peptide deformylase 1bs6. The structure of peptide deformylase was cleaned of unnecessary small molecules and converted to an ICM object using an object conversion protocol that incorporates addition of hydrogens, optimization of His, Asn and Gln rotamers, and assignment of protonation states and secondary structure^95^.

The models of chains A (enzyme) and D (peptide MAS) were prepared by global optimization of side chains and backbone annealing in ICM-Pro. Waters W1 and W2, as defined by ^48^, (waters 2041 and 2042, 1bs6), and Ni^2+^ were maintained in the active site for peptide binding stability calculations. Met1 was mutated to all side chains (except Pro) and relative protein:protein binding free energy ΔΔG calculated for all mutants^97^. Formylated peptides fMAS, fVAS and fAAS were built using the ICM-Pro ligand editor and energy minimised^98^. Formylated peptides were docked into the enzyme active site of 1bs6 chain A including W1 and Ni^2+^ using flexible ligand docking with relaxed covalent geometry, automatic assignment of charged groups, and fixed peptide-like amide bonds. The binding site was defined based on the position of the bound MAS peptide (1bs6, chain D) in the original structure (1bs6, chain A). Docking scores were calculated in ICM-Pro with docking effort of 5 and the top ten conformers stored for analysis of optimal binding poses^95^. 2D interaction diagrams were prepared in ICM-Pro and structural model figures were prepared with Chimera^99^.

### Small RNA isolation

Biological quadruplicates of single colony BL21(DE3)pLysS cells harboring the pULTRA-*tac*-*Empty* or pULTRA-*tac-metY(AAC)* plasmids, and one of the pCOLA*-T7-Empty*, pCOLA-*T7-ValRS*, or pCOLA*-T7-ValRS/fmt* plasmids were used to inoculate 1 mL of LB^M^ supplemented with spectinomycin, chloramphenicol, carbenicillin, or kanamycin as needed and incubated at 37°C overnight with constant shaking. Overnight cultures were diluted 1:100 in 5 mL LB^M^ with appropriate antibiotics and incubated at 37°C for 2 hours, followed by induction with 1 mM of IPTG and a final outgrowth step for an additional 4 hours to allow for sufficient tRNA expression. The cell cultures were pelleted at 5,000 xg for 5 min. The supernatant was discarded, and the cell pellets were snap frozen and stored at −80°C. We adapted the tRNA-seq sample preparation as previously described ^100^ with minor modifications. Briefly, total RNA was extracted by lysing samples in 700 µL QIAzol Lysis reagent (no. 79306, Qiagen). To lyse the cells in QIAzol Lysis reagent, the samples were snap frozen for 5 minutes in dry ice and subsequently thawed for 5 min at room temperature. Next, 150 µL chloroform was added, followed by vortexing and centrifuging at 15,000 x *g* for 15 min at 4 °C to phase separate total RNA from cellular DNA and proteins. The aqueous layer was transferred to a new Eppendorf tube and mixed with 350 µL 70% ethanol containing 50 mM sodium acetate pH=4.5. Larger RNA molecules were separated via a RNeasy MinElute spin column (no. 74204, Qiagen). The flow-through containing the small RNA molecules (< 200 nt) were mixed with 450 µL of 100% ethanol. The small RNA molecules were isolated on a new RNeasy MinElute spin column. The column was washed twice with 500 µL 80% ethanol containing 50 mM sodium acetate pH=4.5. Finally, the small RNA molecules were eluted in 50 mM sodium acetate pH=4.5, 1mM EDTA pH=8.0. Small RNA concentration was determined on the NanoDrop and samples were frozen at −80°C in single-use aliquots (∼ 10 μg).

### RNA oxidation and β-elimination to determine aminoacylation levels

To measure tRNA aminoacylation levels, we performed RNA oxidation and β-elimination, as previously described^54,55^, with minor modifications. Briefly, 10 μg of small RNA was oxidized by the addition of freshly prepared NaIO_4_ to a final concentration of 50 mM in a 20 µL reaction volume for 30 min at 22°C. The reaction was quenched by addition of 2.2 µL 1 M glucose for 5 min at 22°C. Oxidated small RNA was recovered using the Micro Bio Spin P30 columns (BioRad, 7326223) to buffer exchange, and the RNA Clean and Concentrator kit (Zymo, R1018) to purify RNA. Small RNA was eluted in 20 µL RNase-free water and β-elimination was performed by addition of 30 µL 100 mM sodium borate pH = 9.5 (freshly prepared) for 90 min at 45°C. Small RNA was recovered using the Micro Bio Spin P30 columns (BioRad, 7326223) for buffer exchange and the RNA Clean and Concentrator kit (Zymo, R1018) to purify RNA. Small RNA concentration was determined on the NanoDrop and samples were frozen at −80°C in single-use aliquots (∼150 ng).

### tRNAseq library preparation and sequencing

We prepared small RNA cDNA libraries by using the TGIRT-III template-switching reverse transcriptase (InGex, TGIRT50), as previously described^100,101^, with minor modifications. Briefly, all reactions contained 1 µL (∼ 150 ng) treated small RNA sample, 2 µL of 1 μM pre-annealed TGIRT DNA–RNA heteroduplex (prepared by hybridizing equimolar amounts of rArArUrUrGrArGrCrCrUrArArUrGrCrCrUrGrArArArGrArUrCrGrGrArArGrArGrCrArCrArCrGrUr CrU and AGACGTGTGCTCTTCCGATCTTTCAGGCATTAGGCTCAATTN oligonucleotides in RNAse-free H_2_O), 4 µL 5× TGIRT reaction buffer (2.25 M NaCl, 25 mM MgCl2, 100 mM Tris-HCl, pH 7.5), 2 µL of 100 mM DTT, 9 µL RNase-free water and 1 µL TGIRT-III, and incubated at room temperature for 30 min to initiate template-switching. Next, 1 µL of 25 mM dNTPs (Thermo Fisher Scientific) was added to the reaction mixture, and samples were incubated at 60 °C for 1 hour to carry out reverse transcription. RNA was then hydrolyzed by NaOH, and then neutralized by HCl, and the cDNA library was purified using a MinElute PCR purification kit. The resulting cDNA library was ligated to a pre-adenylated DNA adapter, (/5Phos/GATCNNNAGATCGGAAGAGCGTCGTGT/3SpC3/, in which NNN denotes an N, NN or NNN spacer to increase library diversity during sequencing). The pre-adenylated oligonucleotides were prepared by a 5′ DNA adenylation kit (E2610L, NEB) using thermostable 5′ App DNA/RNA ligase (M0319L, NEB) following the manufacturer’s instructions. The cDNA library was purified using a modified SPRI-based single stranded DNA cleanup as previously described^102^. SPRISelect beads (Beckman Coulter, B23317) and 100% isopropanol were added to the DNA adaptor ligated cDNA library reaction to maintain a final concentration of 7.5% PEG and 38% isopropanol. Next, the bound DNA was washed thrice with 70% ethanol and eluted in 20 µL RNAse-free H_2_O. The DNA adaptor ligated cDNA library was amplified and barcoded (compatible Illumina Truseq DNA single index) using Phusion High-Fidelity DNA Polymerase (NEB, M0530S). PCR products were size selected to remove adapter dimers below 200 bp using SPRISelect beads (Beckman Coulter, B23317) according to manufacturer’s instructions. The resultant TruSeq single index barcoded libraries were quantified via NEBNext® Library Quant Kit for Illumina® (NEB, E7630S) and pooled for next-generation sequencing. The pooled libraries were submitted to the Harvard Medical School Biopolymers Facility to be sequenced for 2 x 150 cycles on an Illumina NextSeq platform.

### tRNAseq data analysis

Sequencing libraries were demultiplexed using bcl2fastq (v2.20, Illumina). Following demultiplexing, forward and reverse paired reads were merged using pear (v0.9.6)^103^ and further trimmed using cutadapt (v4.4)^104^ to remove 5’ and 3’ adaptor sequences introduced during the tRNA library preparation. Example commands for merged paired reads and adaptor trimming:

pear -v 10 -m 150 -q 30 -u 0 -j 20 -f “{_sampleName}_R1.fastq.gz" -r "${_sampleName}_R2.fastq.gz" -o "${_sampleName}"

cutadapt -g GATC -a AATTGAGCCTAATGCCTGAA -j 20 -m 10 -q 30 -o "${_sampleName}.fastq" "${_sampleName}.assembled.fastq"

Quantitation and analysis of tRNA expression, charging and modifications were performed using the mim-tRNA-seq computational pipeline (https://github.com/nedialkova-lab/mim-tRNAseq, v1.3.3)^54,55^. Custom input files were generated following the instructions provided in Behrens and Nedialkova^54^. Briefly, the i-tRNA-AAC mutant tRNA sequence and associated intron information (acquired via tRNAScan-SE 2.0^105^) was manually added to *E. coli* .fa and .out files downloaded from tRNAScan-SE database^58^. This generated custom input files: tRNA reference sequence fasta file (mimtRNAseq-PJ35-tRNAome.fa) and corresponding intron information file (mimtRNAseq-PJ35-tRNAome.out) used in the analysis. The following parameters were used:

Custom-t mimtRNAseq-PJ35-tRNAome.fa -o mimtRNAseq-PJ35-tRNAome.out --cluster-id 0.97 --min-cov 0.0005 --max-mismatches 0.1 --control-condition Empty_vector --max-multi 4 --remap --remap-mismatches 0.075 --posttrans-mod-off

Data wrangling and visualization were performed using R packages tidyverse and ggplot2^106^ respectively. Two files generated via the mim-tRNAseq analysis: CCAcounts.csv and mismatchTable.csv were used for further analysis. Initially, tRNA isodecoders not present in 3 out of 4 replicates and below 100 instances (coverage > 100) were filtered out of the dataset. For aminoacylation analysis, the proportion/percentage of complete 3’ CCA ends vs total reads for each tRNA isodecoder represents the charged/aminoacylation levels of the respective tRNA isodecoder. For modification analysis, RT misincorporation rate at each position of an isodecoder is a proxy for the presence of a tRNA modification. Isodecoder tRNA positions with less than 10% were filtered out to detect unambiguous RT signatures for a tRNA modification. Supplementary table S4 describes tRNA positions with significant RT misincorporation and the predicted modification. All statistical analyses were performed using the R package ggpubR^107^ and are found in the text and figure legends. Significant differences are abbreviated as follows: ns (P > 0.05), * (P ≤ 0.05), ** (P ≤ 0.01), *** (P ≤ 0.001), and **** (P ≤ 0.0001).

## Resource availability

### Lead Contact

Requests for further information and resources should be directed to and will be fulfilled by the lead contact, Paul R. Jaschke.

## Materials availability

All unique/stable reagents generated in this study are available from the lead contact without restriction.

## Data and code availability

- The tRNA sequencing data generated in this study have been deposited in the ArrayExpress database under accession number E-MTAB-15868. The dataset includes raw sequencing reads and associated metadata. The mass spectrometry data have been deposited into the MassIVE (https://massive.ucsd.edu/) data repository under the accession number MSV000099552 and are publicly available as of the date of publication.
- This paper does not report original code.
- Any additional information required to reanalyze the data reported in this paper is available from the lead contact upon request.

## Acknowledgements

We thank Ariel Hecht for providing the R code used to produce several figures and Daniel McDougal, Rhys Tuohy, and Pamela Tsoumbris for helpful feedback on the manuscript. We also thank Ardeshir Amirkhani and Gene Hart-Smith for helpful discussions. Some of the research described herein was facilitated by access to the Australian Proteome Analysis Facility (APAF) funded under the Australian Government’s National Collaborative Research Infrastructure Strategy (NCRIS)/Education Investment Fund. DS was supported by a Macquarie Research Excellence PhD Scholarship, AH was supported by a Macquarie Research Excellence PhD Scholarship and CSIRO SynBio FSP Top-Up scholarship, FW was supported by the Ramsay Fellowship of Applied Science, University of Adelaide, and PRJ was supported by the Molecular Sciences Department, Faculty of Science & Engineering, and the Deputy Vice-Chancellor (Research) of Macquarie University. RMV was supported by US Department of Energy (DOE) grant DE-FG02-02ER63445, and NSF Award 2123243 (both to GMC).

## Author Contributions

**Dominic Scopelliti**: Conceptualization, Experimentation, Formal Analysis, Investigation, Methodology, Visualization, Writing - Original Draft, Writing - Review.

**Russel M. Vincent**: Conceptualization, Experimentation, Formal Analysis, Investigation, Methodology, Visualization, Writing - Original Draft, Writing - Review.

**Andras Hutvagner**: Conceptualization, Experimentation, Formal Analysis, Investigation, Methodology, Visualization, Writing - Original Draft, Funding acquisition.

**Fiona Whelan:** Conceptualization, Formal Analysis, Investigation, Methodology, Visualization, Writing - Original Draft, Writing - Review.

**Will Klare:** Formal analysis, Visualization.

**Ignatius Pang:** Formal analysis, Visualization.

**George M. Church**: Writing - Review & Editing, Funding acquisition.

**Paul R. Jaschke**: Conceptualization, Writing - Original Draft, Writing - Review & Editing, Supervision, Project administration, Funding acquisition.

## References

1. 1. de la Torre, D., and Chin, J.W. (2021). Reprogramming the genetic code. Nat Rev Genet 22, 169–184. 10.1038/s41576-020-00307-7.

2. Mukai, T., Lajoie, M.J., Englert, M., and Soll, D. (2017). Rewriting the Genetic Code. Annu Rev Microbiol 71, 557–577. 10.1146/annurev-micro-090816-093247.

3. Reynolds, N.M., Vargas-Rodriguez, O., Soll, D., and Crnkovic, A. (2017). The central role of tRNA in genetic code expansion. Biochim Biophys Acta Gen Subj 1861, 3001–3008. 10.1016/j.bbagen.2017.03.012.

4. Chin, J.W. (2017). Expanding and reprogramming the genetic code. Nature 550, 53–60.

5. Liu, C.C., Jewett, M.C., Chin, J.W., and Voigt, C.A. (2018). Toward an orthogonal central dogma. Nat Chem Biol 14, 103–106. 10.1038/nchembio.2554.

6. Neumann, H. (2012). Rewiring translation - Genetic code expansion and its applications. FEBS Lett 586, 2057–2064. 10.1016/j.febslet.2012.02.002.

7. Amiram, M., Haimovich, A.D., Fan, C., Wang, Y.S., Aerni, H.R., Ntai, I., Moonan, D.W., Ma, N.J., Rovner, A.J., Hong, S.H., et al. (2015). Evolution of translation machinery in recoded bacteria enables multi-site incorporation of nonstandard amino acids. Nat Biotechnol 33, 1272–1279. 10.1038/nbt.3372.

8. Kunjapur, A.M., Stork, D.A., Kuru, E., Vargas-Rodriguez, O., Landon, M., Söll, D., and Church, G.M. (2018). Engineering posttranslational proofreading to discriminate nonstandard amino acids. Proceedings of the National Academy of Sciences 115, 619–624.

9. Link, A.J., Mock, M.L., and Tirrell, D.A. (2003). Non-canonical amino acids in protein engineering. Curr Opin Biotechnol 14, 603–609. 10.1016/j.copbio.2003.10.011.

10. Ma, N.J., and Isaacs, F.J. (2016). Genomic Recoding Broadly Obstructs the Propagation of Horizontally Transferred Genetic Elements. Cell Syst 3, 199–207. 10.1016/j.cels.2016.06.009.

11. Hankore, E.D., Zhang, L., Chen, Y., Liu, K., Niu, W., and Guo, J. (2019). Genetic Incorporation of Noncanonical Amino Acids Using Two Mutually Orthogonal Quadruplet Codons. ACS Synth Biol 8, 1168–1174. 10.1021/acssynbio.9b00051.

12. Aleksashin, N.A., Szal, T., d’Aquino, A.E., Jewett, M.C., Vazquez-Laslop, N., and Mankin, A.S. (2020). A fully orthogonal system for protein synthesis in bacterial cells. Nat Commun 11, 1858. 10.1038/s41467-020-15756-1.

13. Diercks, C.S., Dik, D.A., and Schultz, P.G. (2021). Adding New Chemistries to the Central Dogma of Molecular Biology. Chem 7, 2883–2895. 10.1016/j.chempr.2021.09.014.

14. Kolber, N.S., Fattal, R., Bratulic, S., Carver, G.D., and Badran, A.H. (2021). Orthogonal translation enables heterologous ribosome engineering in E. coli. Nat Commun 12, 599. 10.1038/s41467-020-20759-z.

15. Costello, A., and Badran, A.H. (2021). Synthetic Biological Circuits within an Orthogonal Central Dogma. Trends Biotechnol 39, 59–71. 10.1016/j.tibtech.2020.05.013.

16. Wu, K.L., Moore, J.A., Miller, M.D., Chen, Y., Lee, C., Xu, W., Peng, Z., Duan, Q., Phillips, G.N., Jr., Uribe, R.A., and Xiao, H. (2022). Expanding the eukaryotic genetic code with a biosynthesized 21st amino acid. Protein Sci 31, e4443. 10.1002/pro.4443.

17. Tsoumbris, P.R., Vincent, R.M., and Jaschke, P.R. (2024). Designing a simple and efficient phage biocontainment system using the amber suppressor initiator tRNA. Archives of Virology 169, 248.

18. Schmied, W.H., Tnimov, Z., Uttamapinant, C., Rae, C.D., Fried, S.D., and Chin, J.W. (2018). Controlling orthogonal ribosome subunit interactions enables evolution of new function. Nature 564, 444–448. 10.1038/s41586-018-0773-z.

19. Tharp, J.M., Krahn, N., Varshney, U., and Soll, D. (2020). Hijacking Translation Initiation for Synthetic Biology. Chembiochem 21, 1387–1396. 10.1002/cbic.202000017.

20. Tharp, J.M., Ad, O., Amikura, K., Ward, F.R., Garcia, E.M., Cate, J.H.D., Schepartz, A., and Soll, D. (2020). Initiation of Protein Synthesis with Non-Canonical Amino Acids In Vivo. Angew Chem Int Ed Engl 59, 3122–3126. 10.1002/anie.201914671.

21. Tharp, J.M., Vargas-Rodriguez, O., Schepartz, A., and Soll, D. (2021). Genetic encoding of three distinct noncanonical amino acids using reprogrammed initiator and nonsense codons. ACS chemical biology 16, 766–774.

22. Vincent, R.M., Wright, B.W., and Jaschke, P.R. (2019). Measuring Amber Initiator tRNA Orthogonality in a Genomically Recoded Organism. ACS Synth Biol 8, 675–685. 10.1021/acssynbio.9b00021.

23. Vincent, R., Yiasemides, P., and Jaschke, P. (2019). An orthogonal amber initiator tRNA functions similarly across diverse Escherichia coli laboratory strains. Matters 2019, 1–11.

24. Shetty, S., Shah, R.A., Chembazhi, U.V., Sah, S., and Varshney, U. (2017). Two highly conserved features of bacterial initiator tRNAs license them to pass through distinct checkpoints in translation initiation. Nucleic Acids Res 45, 2040–2050. 10.1093/nar/gkw854.

25. Shah, R.A., Varada, R., Sah, S., Shetty, S., Lahry, K., Singh, S., and Varshney, U. (2019). Rapid formylation of the cellular initiator tRNA population makes a crucial contribution to its exclusive participation at the step of initiation. Nucleic Acids Res 47, 1908–1919. 10.1093/nar/gky1310.

26. Rodnina, M.V. (2018). Translation in Prokaryotes. Cold Spring Harb Perspect Biol 10. 10.1101/cshperspect.a032664.

27. Lahry, K., Datta, M., and Varshney, U. (2024). Genetic analysis of translation initiation in bacteria: An initiator tRNA-centric view. Mol Microbiol. 10.1111/mmi.15243.

28. Shetty, S., and Varshney, U. (2016). An evolutionarily conserved element in initiator tRNAs prompts ultimate steps in ribosome maturation. Proc Natl Acad Sci U S A 113, E6126–E6134. 10.1073/pnas.1609550113.

29. Guillon, J.-M., Meinnel, T., Mechulam, Y., Lazennec, C., Blanquet, S., and Fayat, G. (1992). Nucleotides of tRNA governing the specificity of Escherichia coli methionyl-tRNAfMet formyltransferase. Journal of Molecular Biology 224, 359–367. 10.1016/0022-2836(92)91000-F.

30. Schulman, L.H., and Pelka, H. (1985). In vitro conversion of a methionine to a glutamine-acceptor tRNA. Biochemistry 24, 7309–7314.

31. Mayer, C., Kohrer, C., Kenny, E., Prusko, C., and RajBhandary, U.L. (2003). Anticodon sequence mutants of Escherichia coli initiator tRNA: effects of overproduction of aminoacyl-tRNA synthetases, methionyl-tRNA formyltransferase, and initiation factor 2 on activity in initiation. Biochemistry 42, 4787–4799. 10.1021/bi034011r.

32. Chattapadhyay, R., Pelka, H., and Schulman, L.H. (1990). Initiation of in vivo protein synthesis with non-methionine amino acids. Biochemistry 29, 4263–4268. 10.1021/bi00470a001.

33. Giege, R., and Eriani, G. (2023). The tRNA identity landscape for aminoacylation and beyond. Nucleic Acids Res 51, 1528–1570. 10.1093/nar/gkad007.

34. Borkowski, O., Ceroni, F., Stan, G.-B., and Ellis, T. (2016). Overloaded and stressed: whole-cell considerations for bacterial synthetic biology. Current opinion in microbiology 33, 123–130.

35. Mohler, K., Moen, J.M., Rogulina, S., and Rinehart, J. (2023). System-wide optimization of an orthogonal translation system with enhanced biological tolerance. Molecular Systems Biology 19, e10591.

36. Abbott, J.A., Francklyn, C.S., and Robey-Bond, S.M. (2014). Transfer RNA and human disease. Front Genet 5, 158. 10.3389/fgene.2014.00158.

37. Hecht, A., Glasgow, J., Jaschke, P.R., Bawazer, L.A., Munson, M.S., Cochran, J.R., Endy, D., and Salit, M. (2017). Measurements of translation initiation from all 64 codons in E. coli. Nucleic acids research 45, 3615–3626.

38. Love, M.I., Huber, W., and Anders, S. (2014). Moderated estimation of fold change and dispersion for RNA-seq data with DESeq2. Genome biology 15, 550.

39. Mertens, N., Remaut, E., and Fiers, W. (1995). Tight transcriptional control mechanism ensures stable high-level expression from T7 promoter-based expression plasmids. Bio/Technology 13, 175–179.

40. Brophy, J.A., and Voigt, C.A. (2014). Principles of genetic circuit design. Nature methods 11, 508–520.

41. Varshney, U., and RajBhandary, U.L. (1990). Initiation of protein synthesis from a termination codon. Proc Natl Acad Sci U S A 87, 1586–1590. 10.1073/pnas.87.4.1586.

42. Fukai, S., Nureki, O., Sekine, S., Shimada, A., Vassylyev, D.G., and Yokoyama, S. (2003). Mechanism of molecular interactions for tRNA(Val) recognition by valyl-tRNA synthetase. RNA 9, 100–111. 10.1261/rna.2760703.

43. Horowitz, J., Chu, W.C., Derrick, W.B., Liu, J.C., Liu, M., and Yue, D. (1999). Synthetase recognition determinants of E. coli valine transfer RNA. Biochemistry 38, 7737–7746. 10.1021/bi990490b.

44. Chang, S.Y., McGary, E.C., and Chang, S. (1989). Methionine aminopeptidase gene of Escherichia coli is essential for cell growth. Journal of Bacteriology 171, 4071–4072. 10.1128/jb.171.7.4071-4072.1989.

45. Ben-Bassat, A., Bauer, K., Chang, S.Y., Myambo, K., Boosman, A., and Chang, S. (1987). Processing of the initiation methionine from proteins: properties of the Escherichia coli methionine aminopeptidase and its gene structure. J Bacteriol 169, 751–757. 10.1128/jb.169.2.751-757.1987.

46. Firnberg, E., Labonte, J.W., Gray, J.J., and Ostermeier, M. (2014). A comprehensive, high-resolution map of a gene’s fitness landscape. Molecular biology and evolution 31, 1581–1592.

47. Adams, J.M. (1968). On the release of the formyl group from nascent protein. Journal of Molecular Biology 33, 571–589. 10.1016/0022-2836(68)90307-0.

48. Becker, A., Schlichting, I., Kabsch, W., Groche, D., Schultz, S., and Wagner, A.V. (1998). Iron center, substrate recognition and mechanism of peptide deformylase. Nature structural biology 5, 1053–1058.

49. Wu, X.-H., Quan, J.-M., and Wu, Y.-D. (2007). Theoretical study of the catalytic mechanism and metal-ion dependence of peptide deformylase. The Journal of Physical Chemistry B 111, 6236–6244.

50. Guillon, J.-M., Mechulam, Y., Schmitter, J., Blanquet, S., and Fayat, G. (1992). Disruption of the gene for Met-tRNA (fMet) formyltransferase severely impairs growth of Escherichia coli. Journal of bacteriology 174, 4294–4301.

51. Mazel, D., Pochet, S., and Marliere, P. (1994). Genetic characterization of polypeptide deformylase, a distinctive enzyme of eubacterial translation. The EMBO journal 13, 914–923.

52. Liao, Y.D., Jeng, J.C., Wang, C.F., Wang, S.C., and Chang, S.T. (2004). Removal of N-terminal methionine from recombinant proteins by engineered E. coli methionine aminopeptidase. Protein Science 13, 1802–1810.

53. Evans, M.E., Clark, W.C., Zheng, G., and Pan, T. (2017). Determination of tRNA aminoacylation levels by high-throughput sequencing. Nucleic Acids Research 45, e133–e133. 10.1093/nar/gkx514.

54. Behrens, A., and Nedialkova, D.D. (2022). Experimental and computational workflow for the analysis of tRNA pools from eukaryotic cells by mim-tRNAseq. STAR Protoc 3, 101579. 10.1016/j.xpro.2022.101579.

55. Behrens, A., Rodschinka, G., and Nedialkova, D.D. (2021). High-resolution quantitative profiling of tRNA abundance and modification status in eukaryotes by mim-tRNAseq. Mol Cell 81, 1802–1815 e1807. 10.1016/j.molcel.2021.01.028.

56. De Crécy-Lagard, V., and Jaroch, M. (2021). Functions of Bacterial tRNA Modifications: From Ubiquity to Diversity. Trends in Microbiology 29, 41–53. 10.1016/j.tim.2020.06.010.

57. Takakura, M., Ishiguro, K., Akichika, S., Miyauchi, K., and Suzuki, T. (2019). Biogenesis and functions of aminocarboxypropyluridine in tRNA. Nature communications 10, 5542.

58. Chan, P.P., and Lowe, T.M. (2009). GtRNAdb: a database of transfer RNA genes detected in genomic sequence. Nucleic acids research 37, D93–D97.

59. Traxler, M.F., Summers, S.M., Nguyen, H.T., Zacharia, V.M., Hightower, G.A., Smith, J.T., and Conway, T. (2008). The global, ppGpp-mediated stringent response to amino acid starvation in Escherichia coli. Molecular microbiology 68, 1128–1148.

60. Qian, Y., Huang, H.-H., Jiménez, J.I., and Del Vecchio, D. (2017). Resource Competition Shapes the Response of Genetic Circuits. ACS Synthetic Biology 6, 1263–1272. 10.1021/acssynbio.6b00361.

61. Ceroni, F., Boo, A., Furini, S., Gorochowski, T.E., Borkowski, O., Ladak, Y.N., Awan, A.R., Gilbert, C., Stan, G.-B., and Ellis, T. (2018). Burden-driven feedback control of gene expression. Nature methods 15, 387–393.

62. Govindan, A., Miryala, S., Mondal, S., and Varshney, U. (2018). Development of assay systems for amber codon decoding at the steps of initiation and elongation in mycobacteria. J Bacteriol 200, e00372–00318. 10.1128/JB.00372-18.

63. Varshney, U., and RajBhandary, U.L. (1992). Role of methionine and formylation of initiator tRNA in initiation of protein synthesis in Escherichia coli. J Bacteriol 174, 7819–7826. 10.1128/jb.174.23.7819-7826.1992.

64. Tardif, K.D., and Horowitz, J. (2002). Transfer RNA determinants for translational editing by Escherichia coli valyl-tRNA synthetase. Nucleic Acids Res 30, 2538–2545. 10.1093/nar/30.11.2538.

65. Spinck, M., Guppy, A., and Chin, J.W. (2025). Automated orthogonal tRNA generation. Nature Chemical Biology 21, 657–667.

66. Giege, R., Ebel, J., and Clark, B. (1973). Formylation of mischarged E. coli tRNA Met f. FEBS letters 30, 291–295.

67. Tardif, K.D., Liu, M., Vitseva, O., Hou, Y.M., and Horowitz, J. (2001). Misacylation and editing by Escherichia coli valyl-tRNA synthetase: evidence for two tRNA binding sites. Biochemistry 40, 8118–8125. 10.1021/bi0103213.

68. Raunio, R., and Rosenqvist, H. (1970). Amino acid pool of Escherichia coli during the different phases of growth. Acta Chem. Scand 24.

69. Fersht, A.R., and Kaethner, M.M. (1976). Enzyme hyperspecificity. Rejection of threonine by the valyl-tRNA synthetase by misacylation and hydrolytic editing. Biochemistry 15, 3342–3346.

70. Jakubowski, H., and R. Fersht, A. (1981). Alternative pathways for editing non-cognate amino acids by aminoacyl-tRNA synthetases. Nucleic acids research 9, 3105–3117.

71. Tardif, K.D., and Horowitz, J. (2004). Functional group recognition at the aminoacylation and editing sites of E. coli valyl-tRNA synthetase. RNA 10, 493–503. 10.1261/rna.5166704.

72. Du, B., Yang, L., Lloyd, C.J., Fang, X., and Palsson, B.O. (2019). Genome-scale model of metabolism and gene expression provides a multi-scale description of acid stress responses in Escherichia coli. PLoS Comput Biol 15, e1007525. 10.1371/journal.pcbi.1007525.

73. Ceroni, F., Algar, R., Stan, G.-B., and Ellis, T. (2015). Quantifying cellular capacity identifies gene expression designs with reduced burden. Nature methods 12, 415–418.

74. Costello, A., Peterson, A.A., Chen, P.-H., Bagirzadeh, R., Lanster, D.L., and Badran, A.H. (2024). Genetic code expansion history and modern innovations. Chemical Reviews 124, 11962–12005.

75. Kim, Y., Cho, S., Kim, J.-C., and Park, H.-S. (2024). tRNA engineering strategies for genetic code expansion. Frontiers in Genetics 15, 1373250.

76. Huang, Y., Zhang, P., Wang, H., Chen, Y., Liu, T., and Luo, X. (2024). Genetic Code Expansion: Recent Developments and Emerging Applications. Chemical Reviews 125, 523–598.

77. Nyerges, A., Chiappino-Pepe, A., Budnik, B., Baas-Thomas, M., Flynn, R., Yan, S., Ostrov, N., Liu, M., Wang, M., Zheng, Q., et al. (2024). Synthetic genomes unveil the effects of synonymous recoding. bioRxiv, 2024.2006.2016.599206. 10.1101/2024.06.16.599206.

78. Robertson, W.E., Rehm, F.B.H., Spinck, M., Schumann, R.L., Tian, R., Liu, W., Gu, Y., Kleefeldt, A.A., Day, C.F., Liu, K.C., et al. (2025). *Escherichia coli* with a 57-codon genetic code. Science 390, eady4368. doi:10.1126/science.ady4368.

79. Calles, J., Justice, I., Brinkley, D., Garcia, A., and Endy, D. (2019). Fail-safe genetic codes designed to intrinsically contain engineered organisms. Nucleic Acids Res 47, 10439–10451. 10.1093/nar/gkz745.

80. Debenedictis, E.A., Carver, G.D., Chung, C.Z., Söll, D., and Badran, A.H. (2021). Multiplex suppression of four quadruplet codons via tRNA directed evolution. Nature Communications 12. 10.1038/s41467-021-25948-y.

81. Awawdeh, A., Radecki, A.A., and Vargas-Rodriguez, O. (2024). Suppressor tRNAs at the interface of genetic code expansion and medicine. Frontiers in Genetics 15, 1420331.

82. Meyer, A.J., Segall-Shapiro, T.H., Glassey, E., Zhang, J., and Voigt, C.A. (2019). Escherichia coli "Marionette" strains with 12 highly optimized small-molecule sensors. Nat Chem Biol 15, 196–204. 10.1038/s41589-018-0168-3.

83. Hecht, A., Filliben, J., Munro, S.A., and Salit, M. (2018). A minimum information standard for reproducing bench-scale bacterial cell growth and productivity. Communications biology 1, 1–9.

84. Schultz, K.C., Supekova, L., Ryu, Y., Xie, J., Perera, R., and Schultz, P.G. (2006). A Genetically Encoded Infrared Probe. Journal of the American Chemical Society 128, 13984–13985. 10.1021/ja0636690.

85. Wang, X.C., Wilson, S.C., and Hammond, M.C. (2016). Next-generation RNA-based fluorescent biosensors enable anaerobic detection of cyclic di-GMP. Nucleic Acids Res 44, e139. 10.1093/nar/gkw580.

86. Sprouffske, K., and Wagner, A. (2016). Growthcurver: an R package for obtaining interpretable metrics from microbial growth curves. BMC bioinformatics 17, 172.

87. MacLean, B., Tomazela, D.M., Shulman, N., Chambers, M., Finney, G.L., Frewen, B., Kern, R., Tabb, D.L., Liebler, D.C., and MacCoss, M.J. (2010). Skyline: an open source document editor for creating and analyzing targeted proteomics experiments. Bioinformatics *26*, 966-968.

88. Kong, A.T., Leprevost, F.V., Avtonomov, D.M., Mellacheruvu, D., and Nesvizhskii, A.I. (2017). MSFragger: ultrafast and comprehensive peptide identification in mass spectrometry-based proteomics. Nat Methods 14, 513–520. 10.1038/nmeth.4256.

89. Demichev, V., Messner, C.B., Vernardis, S.I., Lilley, K.S., and Ralser, M. (2020). DIA-NN: neural networks and interference correction enable deep proteome coverage in high throughput. Nature methods 17, 41–44.

90. Ritchie, M.E., Phipson, B., Wu, D., Hu, Y., Law, C.W., Shi, W., and Smyth, G.K. (2015). limma powers differential expression analyses for RNA-sequencing and microarray studies. Nucleic acids research 43, e47–e47.

91. Molania, R., Gagnon-Bartsch, J.A., Dobrovic, A., and Speed, T.P. (2019). A new normalization for Nanostring nCounter gene expression data. Nucleic acids research 47, 6073–6083.

92. Poulos, R.C., Hains, P.G., Shah, R., Lucas, N., Xavier, D., Manda, S.S., Anees, A., Koh, J.M., Mahboob, S., and Wittman, M. (2020). Strategies to enable large-scale proteomics for reproducible research. Nature communications 11, 3793.

93. Storey, J.D. (2003). The positive false discovery rate: a Bayesian interpretation and the q-value. The annals of statistics 31, 2013–2035.

94. Yu, G., Wang, L.-G., Han, Y., and He, Q.-Y. (2012). clusterProfiler: an R package for comparing biological themes among gene clusters. Omics: a journal of integrative biology 16, 284–287.

95. Abagyan, R., Totrov, M., and Kuznetsov, D. (1994). ICM—A new method for protein modeling and design: Applications to docking and structure prediction from the distorted native conformation. Journal of computational chemistry 15, 488–506.

96. Joosten, R.P., Long, F., Murshudov, G.N., and Perrakis, A. (2014). The PDB_REDO server for macromolecular structure model optimization. IUCrJ 1, 213–220.

97. Schapira, M., Totrov, M., and Abagyan, R. (1999). Prediction of the binding energy for small molecules, peptides and proteins. Journal of Molecular Recognition 12, 177–190.

98. Totrov, M., and Abagyan, R. (1997). Flexible protein–ligand docking by global energy optimization in internal coordinates. Proteins: Structure, Function, and Bioinformatics 29, 215–220.

99. Pettersen, E.F., Goddard, T.D., Huang, C.C., Couch, G.S., Greenblatt, D.M., Meng, E.C., and Ferrin, T.E. (2004). UCSF Chimera—a visualization system for exploratory research and analysis. Journal of computational chemistry 25, 1605–1612.

100. Nyerges, A., Vinke, S., Flynn, R., Owen, S.V., Rand, E.A., Budnik, B., Keen, E., Narasimhan, K., Marchand, J.A., and Baas-Thomas, M. (2023). A swapped genetic code prevents viral infections and gene transfer. Nature 615, 720–727.

101. Xu, H., Yao, J., Wu, D.C., and Lambowitz, A.M. (2019). Improved TGIRT-seq methods for comprehensive transcriptome profiling with decreased adapter dimer formation and bias correction. Scientific reports 9, 7953.

102. Fishman, A., Light, D., and Lamm, A.T. (2018). QsRNA-seq: a method for high-throughput profiling and quantifying small RNAs. Genome Biology 19, 113. 10.1186/s13059-018-1495-0.

103. Zhang, J., Kobert, K., Flouri, T., and Stamatakis, A. (2013). PEAR: a fast and accurate Illumina Paired-End reAd mergeR. Bioinformatics 30, 614–620. 10.1093/bioinformatics/btt593.

104. Martin, M. (2011). Cutadapt removes adapter sequences from high-throughput sequencing reads. 2011 *17*, 3. 10.14806/ej.17.1.200.

105. Chan, Patricia P., Lin, Brian Y., Mak, Allysia J., and Lowe, Todd M. (2021). tRNAscan-SE 2.0: improved detection and functional classification of transfer RNA genes. Nucleic Acids Research 49, 9077–9096. 10.1093/nar/gkab688.

106. Wickham, H. (2016). ggplot2: elegant graphics for data analysis (Springer International Publishing). 10.1007/978-3-319-24277-4.

107. Kassambara, A. (2023). ggpubr: ’ggplot2’ Based Publication Ready Plots.

