## Supplementary File for "Defining bottlenecks and physiological impact of an orthogonal translation initiation system"

[3]Present address: Department of Biochemistry & Molecular Biophysics, Washington University School of Medicine, St. Louis, MO, USA

[6]Present address: Macquarie Medical School, Macquarie University, Sydney, NSW, Australia

[10]Lead contact

**CONTENTS**

**Supporting Figures**

| | |
|---|---|
| Figure S1 | Secondary structures of *E. coli* i-tRNA-fMet-CAU-2, mutant i-tRNA-AAC, and e-tRNA-Val-UAC-1. |
| Figure S2 | Bulk fluorescence measurements of a (NNN)sfGFP reporter library in the presence and absence of the mutant i-tRNA-AAC. |
| Figure S3 | Violin Plots of Flow Cytometry Data. |
| Figure S4 | LC-MS/MS detection of N-terminal peptides of (AUG)Nluc reporter protein purified from *E. coli* BL21(DE3)pLysS (pULTRA-*tac-Empty*; pET20b-*T7-(AUG)Nluc6HIS*). |

Figure S5 LC-MS/MS detection of low abundance N-terminal peptides from (AUG)Nluc reporter protein bearing glycine or threonine at the N-terminus purified from *E. coli* BL21(DE3)pLysS (pULTRA-*tac-Empty*; pET20b-*T7-(AUG)Nluc6HIS*).

Figure S6 LC-MS/MS detection of N-terminal peptides of (GUU)Nluc reporter protein purified from *E. coli* BL21(DE3)pLysS (pULTRA-*tac-Empty*; pET20b-*T7*-(*GUU)Nluc6HIS*).

Figure S7 LC-MS/MS detection of N-terminal peptides of (GUU)Nluc reporter protein purified from *E. coli* BL21(DE3)pLysS (pULTRA-*tac-metY*(AAC); pET20b-*T7*-(*GUU)Nluc6HIS*).

Figure S8 LC-MS/MS detection of low abundance N-terminal peptides from (AUG)Nluc reporter protein bearing glycine, alanine, or isoleucine at the N-terminus purified from *E. coli* BL21(DE3)pLysS (pULTRA-*tac-Empty*; pET20b-*T7*-*(AUG)Nluc6HIS*).

Figure S9 Schematic of pCOLA plasmids designed to overexpress ValRS or ValRS/FMT.

Figure S10 Reduction in native (AUG)sfGFP expression and strain fitness during ValRS or ValRS/FMT overexpression.

Figure S11 Differential expression analysis of tRNA measured from *E. coli* BL21(DE3)pLysS cells expressing i-tRNA-AAC with either ValRS, or both ValRS and FMT, compared to cells harbouring an empty pCOLA plasmid**.**

Figure S12 Aminoacylation level of all tRNA isodecoders.

Figure S13 Expression of ValRS or ValRS/FMT influence tRNA modifications that induce reverse transcription (RT) mismatches.

Figure S14 Evaluation of the DIA analysis.

**Supporting Tables**

Table S1 ICM-Pro ligand docking scores for peptides fMAS, fVAS and fAAS

Table S2 ICM-Pro peptide docking scores fMAS

Table S3 ICM-Pro peptide docking scores fVAS

Table S4 tRNA positions from mim-tRNAseq dataset with more than 10% reverse-transcription misincorporation.

Table S5 Gene Sequences

Table S6 Oligos used (sequencing, Gibson assembly, colony PCR for all plasmids)

**Supporting Files**

Supporting File S1: DDA inclusion list

Supporting File S2: Bulk fluorescence data

Supporting File S3: Differential expression analysis

Supporting File S4: Fitness Assay

Supporting File S5: Plasmid sequence: pULTRA-*tac-metY*(AAC)

Supporting File S6: Plasmid sequence: pET20b-*T7-(NNN)sfGFP*

Supporting File S7: Plasmid sequence pET20b-*T7-(GUU)Nluc*

Supporting File S8: Plasmid sequence pET20b-*T7-Nluc*(AUG)

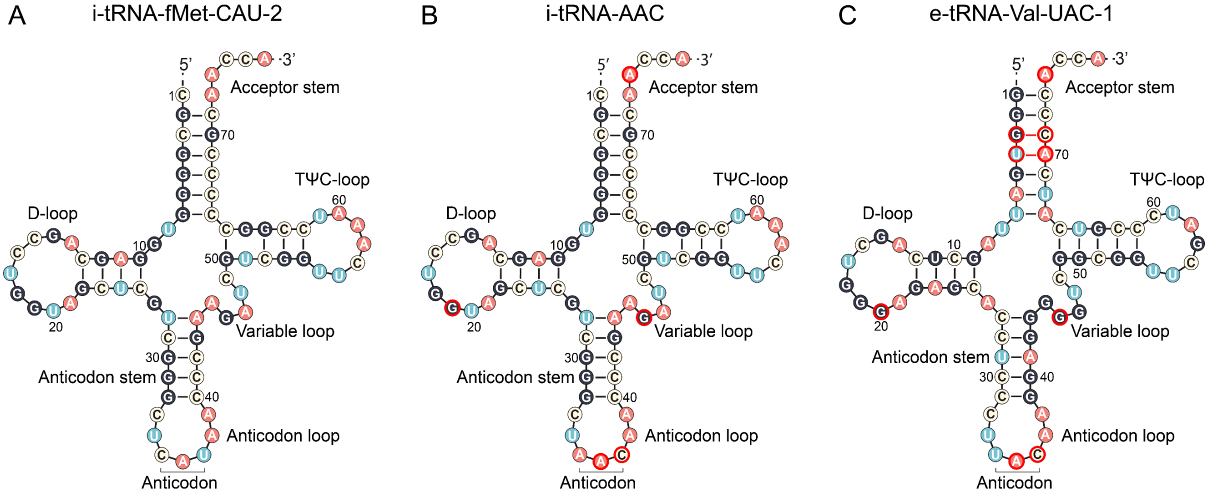


**Figure S1. Secondary structures of *E. coli* i-tRNA-fMet-CAU-2 and mutant i-tRNA-AAC.** (A) *E. coli* initiator tRNA-fMet-CAU-2 sequence retrieved from GtRNAdb database[1]. (B) Sequence of *E. coli* initiator tRNA with AAC anticodon mutation. (C) *E. coli* e-tRNA-Val-UAC-1 sequence. Nucleotides outlined in red highlight the known sequence motifs recognized by valyl-tRNA synthetase.

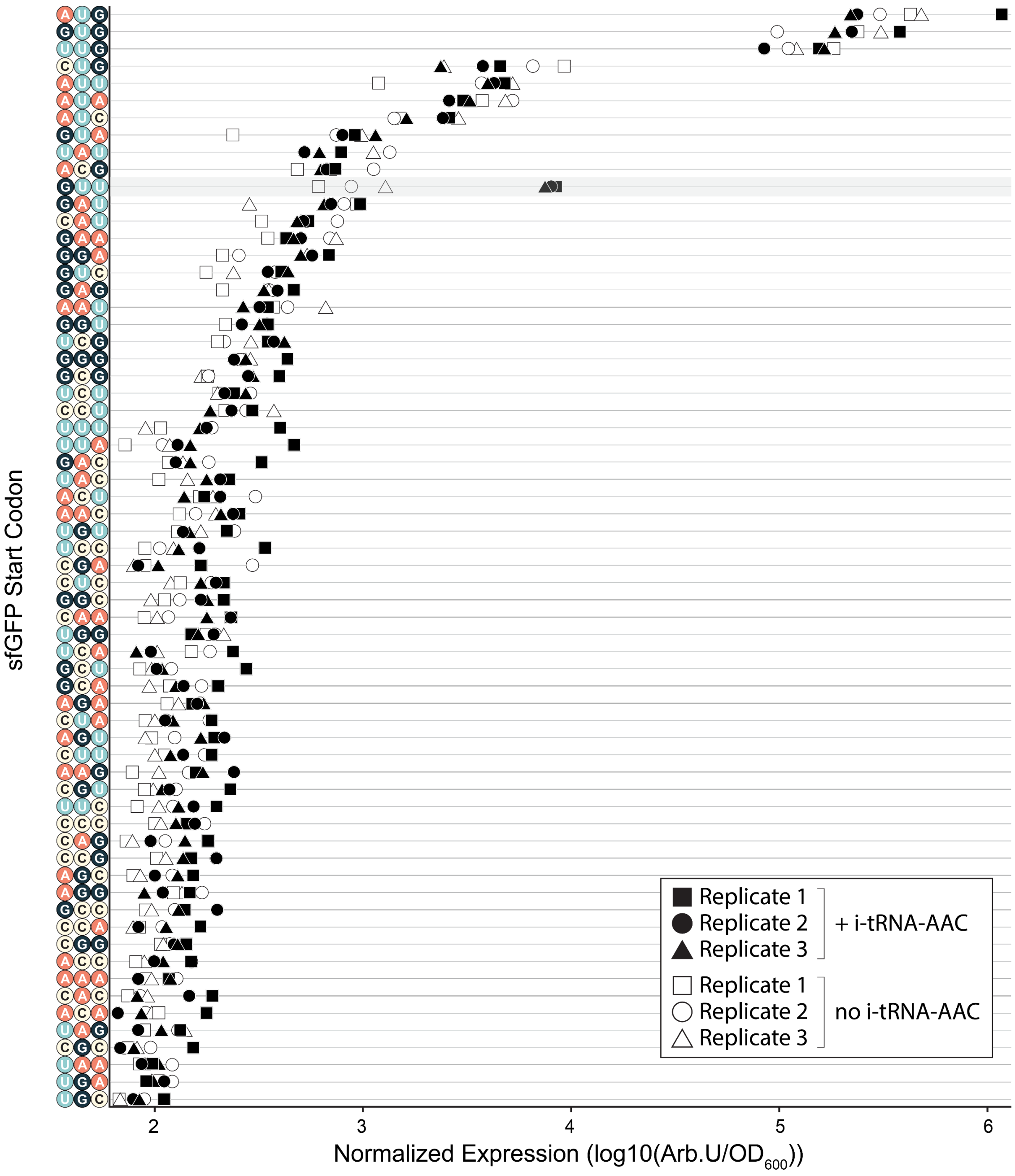


**Figure S2. Bulk fluorescence measurements of a (NNN)sfGFP reporter library in the presence and absence of the mutant i-tRNA-AAC.** Normalized bulk cell fluorescence (Arb. U./$OD_{600}$) of all sfGFP start codon variants were captured from cells grown to mid-log phase (0.6 $OD_{600}$) in the presence of the i-tRNA-AAC induced with 1 mM IPTG for 1-hour prior to measurement. Replicate 1 is represented by squares, replicate 2 by circles, and replicate 3 by triangles. Solid filled data points indicate samples where i-tRNA-AAC was co-expressed, whereas unfilled data points indicate control samples where no mutant i-tRNA was expressed.

A

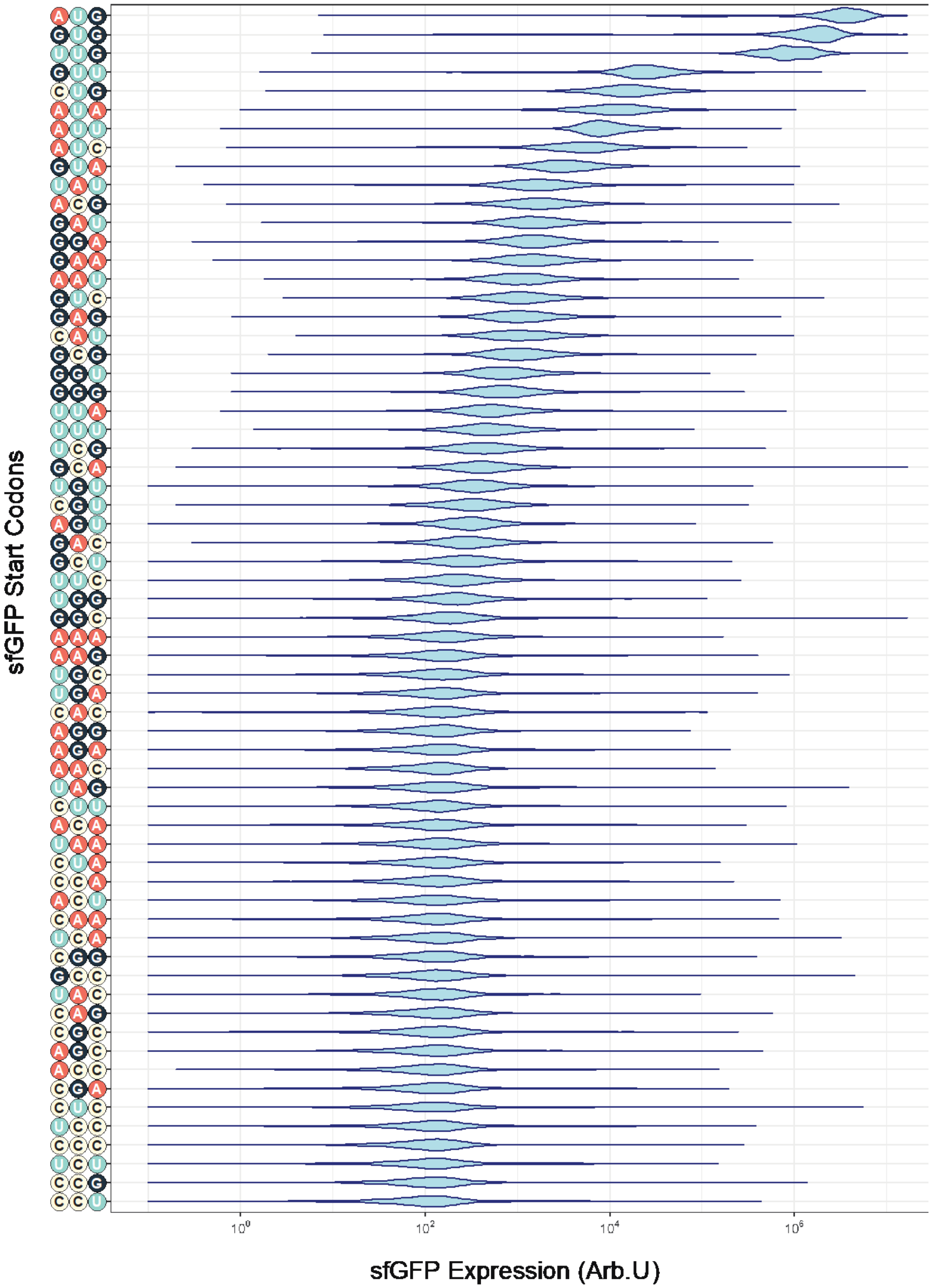
sfGFP Start Codons
AUG
GUG
UUG
GUU
CUG
AUA
AUU
AUC
GUA
UAU
ACG
GAU
GGA
GAA
AAU
GUC
GAG
CAU
GCG
GGU
GGG
UUA
UUU
UCG
GCA
UGU
CGU
AGU
GAC
GCU
UUC
UGG
GGC
AAA
AAG
UGC
UGA
CAC
AGG
AGA
AAC
UAG
CUU
ACA
UAA
CUA
CCA
ACU
CAA
UCA
CGG
GCC
UAC
CAG
CGC
AGC
ACC
CGA
CUC
UCC
CCC
UCU
CCG
CCU
$10^0$
$10^2$
$10^4$
$10^6$
sfGFP Expression (Arb.U)

B

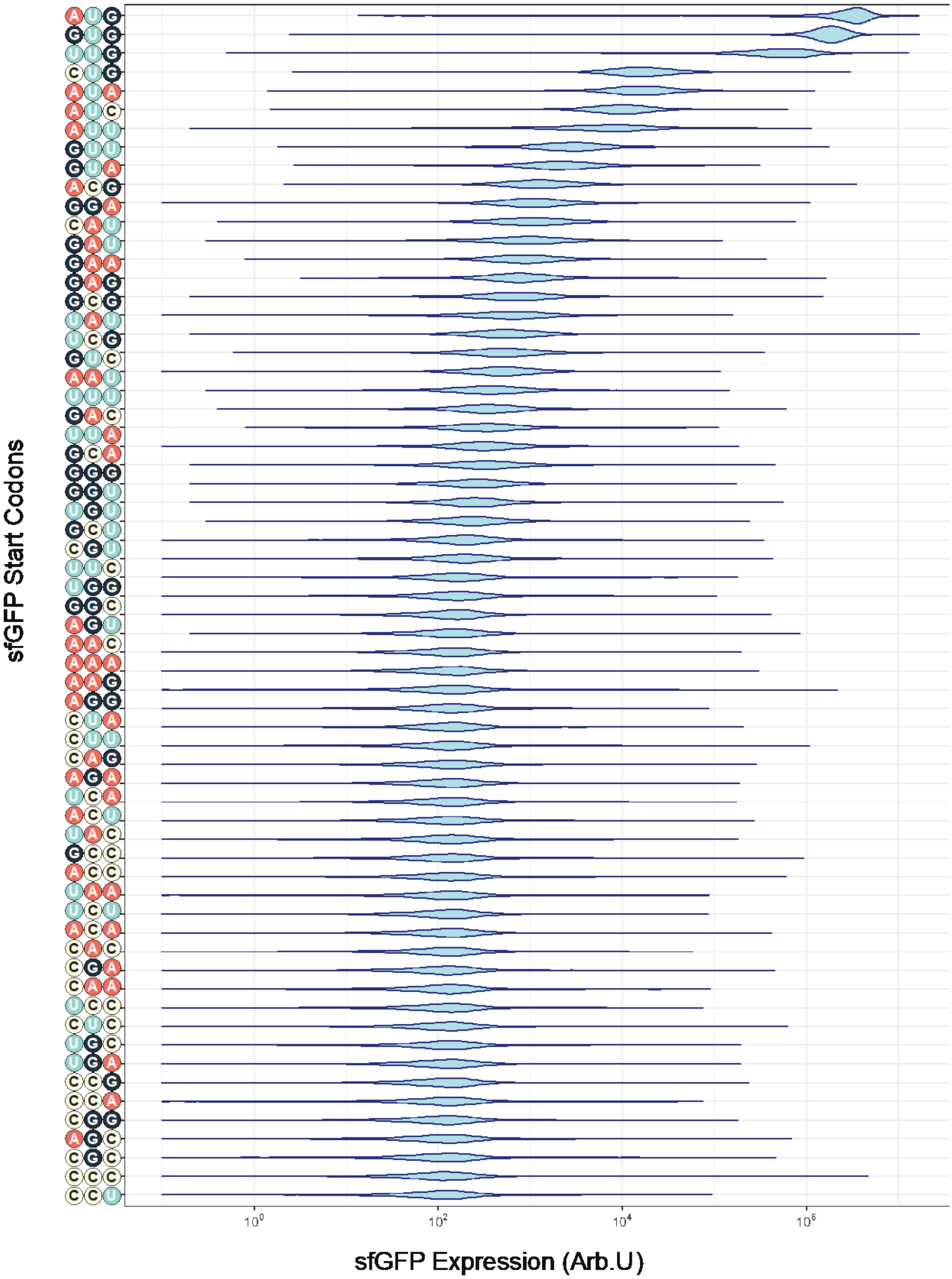


**Figure S3. Violin Plots of Flow Cytometry Data.** Violin plots of the flow cytometry data for both (A) pULTRA-*tac-metY*(AAC) and (B) pULTRA-*tac-Empty* strains expressing sfGFP from all possible 64 start codons. Cells were measured on a CytoFLEX S (Beckman Coulter) using a FITC fluorescence channel with a 488 nm excitation laser and 525/40 nm emission band-pass filter with side scatter triggered events threshold set to 10,000 events.

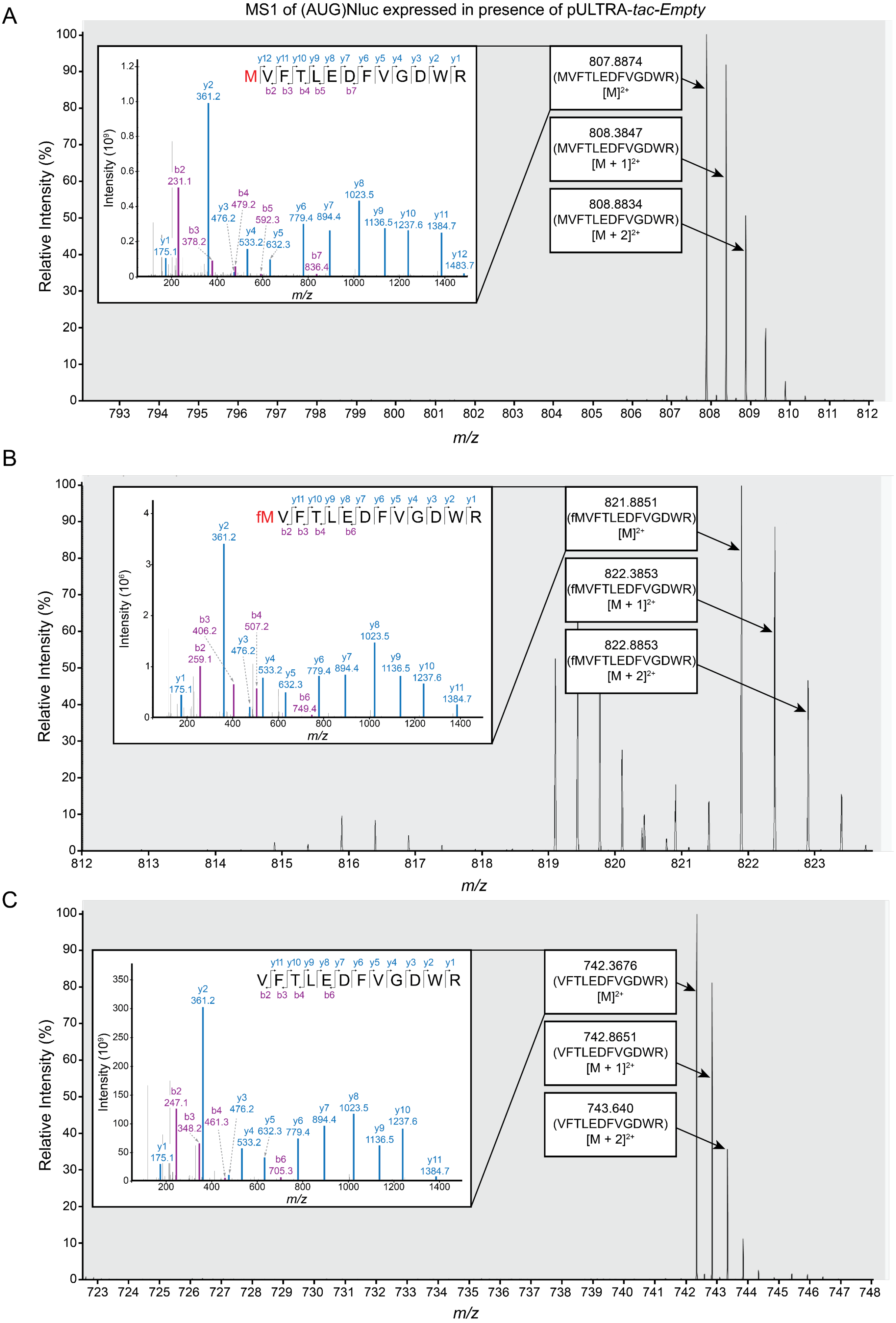


**Figure S4. LC-MS/MS detection of N-terminal peptides of (AUG)Nluc reporter protein purified from *E. coli* BL21(DE3)pLysS (pULTRA-*tac-Empty*; pET20b-*T7-(AUG)Nluc6HIS*).** (A) MS1 scan during retention time range of 45-46 minutes. Labelled monoisotopic peak (807.8874 *m/z*) consistent with the N-terminal peptide bearing methionine (MVFTLEDFVGDWR) and its respective isotopes are shown. The inset shows MS2 scan with $y^-$ and $b^-$ series product ion signals used to confirm amino acid sequence of N-terminal peptide. Schematic of N-terminal peptide (top right of insets) shows peptide fragmentation patterns occurring to produce each of the MS2 ions. (B) MS1 scan during retention time range of 49.5-50 minutes showing monoisotopic peak (821.8851 *m/z*) consistent with the N-terminal peptide bearing formylated methionine (fMVFTLEDFVGDWR) at the N-terminus. MS2 product ions and peptide schematic are shown in inset. (C) MS1 scan during retention time range of 44.5-46 minutes showing the monoisotopic peak (742.3676 *m/z*) consistent with a N-terminal peptide with the residue at N-terminus removed (VFTLEDFVGDWR). MS2 product ions and peptide schematic are also shown in inset.

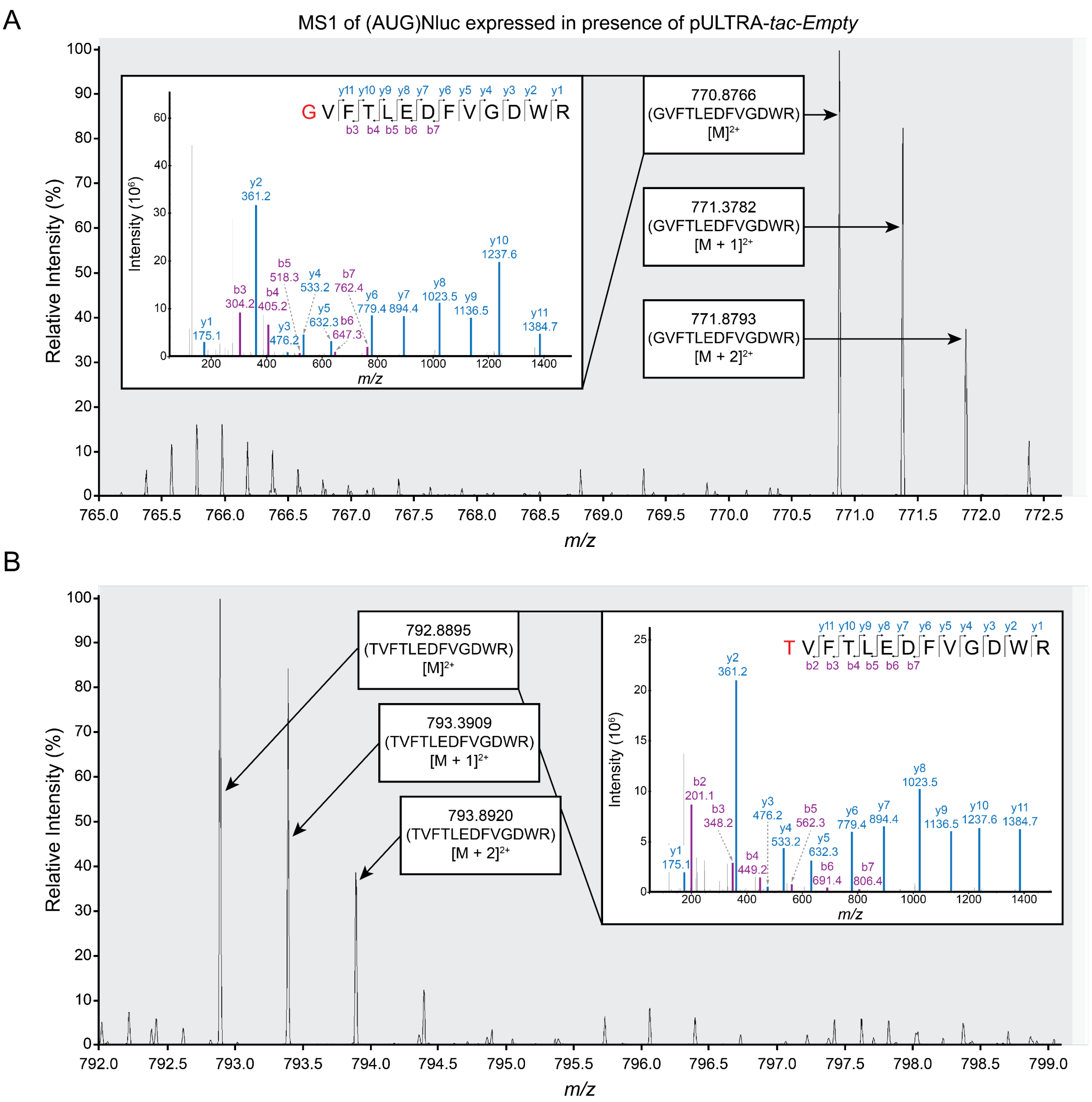


**Figure S5. LC-MS/MS detection of low abundance N-terminal peptides from (AUG)Nluc reporter protein bearing glycine or threonine at the N-terminus purified from *E. coli* BL21(DE3)pLysS (pULTRA-*tac-Empty*; pET20b-*T7-(AUG)Nluc6HIS*).** (A) MS1 scan during retention time range of 44.5-45.5 minutes with labelled monoisotopic peak (770.8766 *m/z*) consistent with an N-terminal peptide bearing glycine (GVFTLEDFVGDWR) and its respective isotopes is shown. The inset shows MS2 scan at with $y^{-}$ and $b^{-}$ series product ion signals used to confirm amino acid sequence of N-terminal peptide. (B) MS1 scan during retention time range of 44-45 with labelled monoisotopic peak (792.8895 *m/z*) consistent with an N-terminal peptide bearing threonine (TVFTLEDFVGDWR) and its respective isotopes. The inset shows MS2 scan with $y^{-}$ and $b^{-}$ series product ion signals used to confirm amino acid sequence of N-terminal peptide. Schematic of N-terminal peptide (top right of insets) shows peptide fragmentation patterns occurring to produce each of the MS2 product ions.

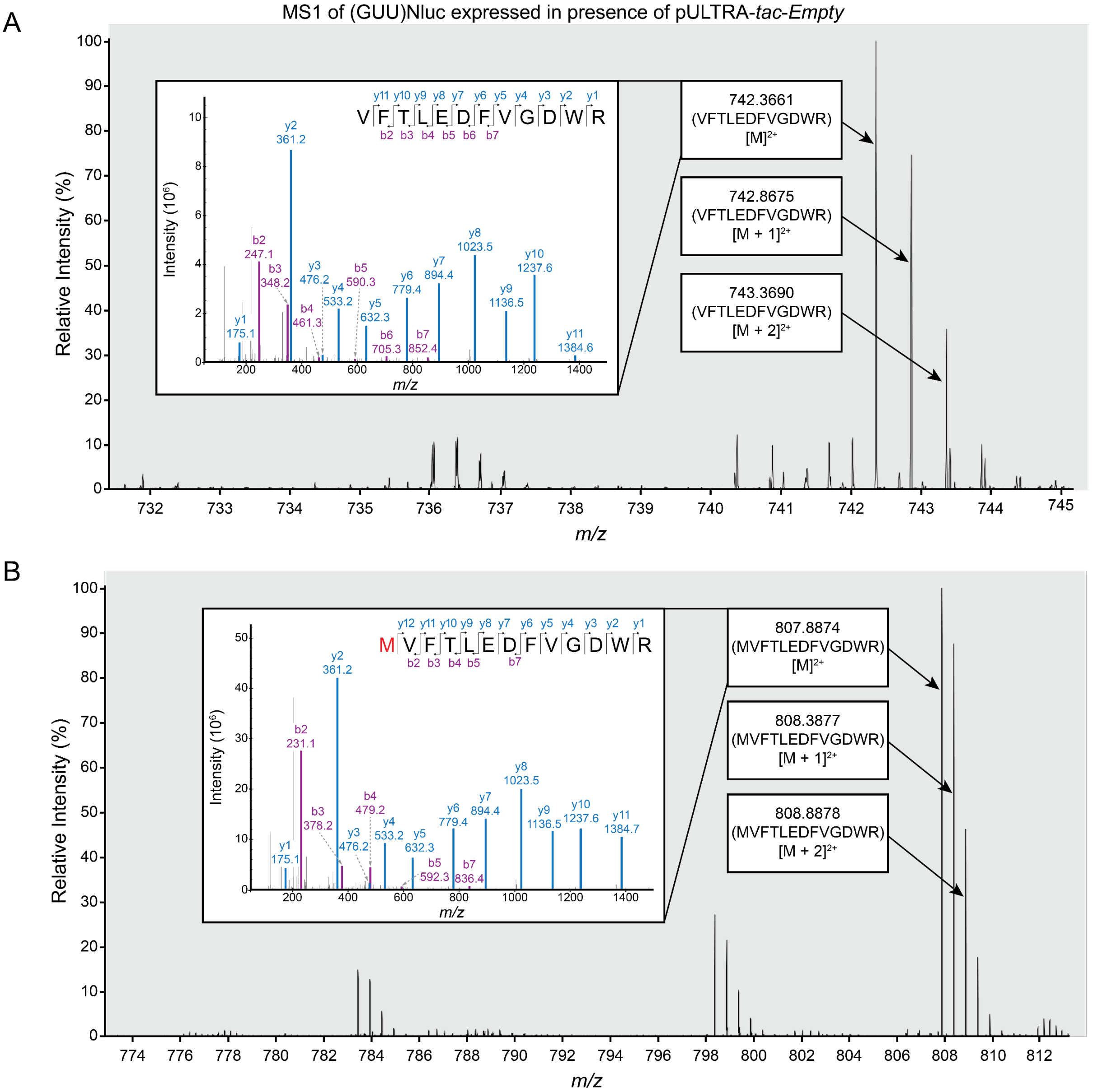


**Figure S6. LC-MS/MS detection of N-terminal peptides of (GUU)Nluc reporter protein purified from *E. coli* BL21(DE3)pLysS (pULTRA-*tac-Empty*; pET20b-*T7*-(*GUU)Nluc6HIS*).** (A) MS1 scan during retention time range of 44.5-46 minutes. Labelled monoisotopic peak (742.3661 *m/z*) consistent with the N-terminal peptide with residue at N-terminus removed (VFTLEDFVGDWR) and its respective isotopes are shown. The inset shows MS2 scan with $y^-$ and $b^-$ series product ion signals used to confirm amino acid sequence of N-terminal peptide. Schematic of N-terminal peptide (top right of insets) shows peptide fragmentation patterns occurring to produce each of the MS2 ions. (B) MS1 scan during retention time range of 46-46.9 minutes showing monoisotopic peak (807.8874 *m/z*) consistent with the N-terminal peptide bearing methionine (MVFTLEDFVGDWR) at the N-terminus. MS2 product ions and peptide schematic are shown in inset.

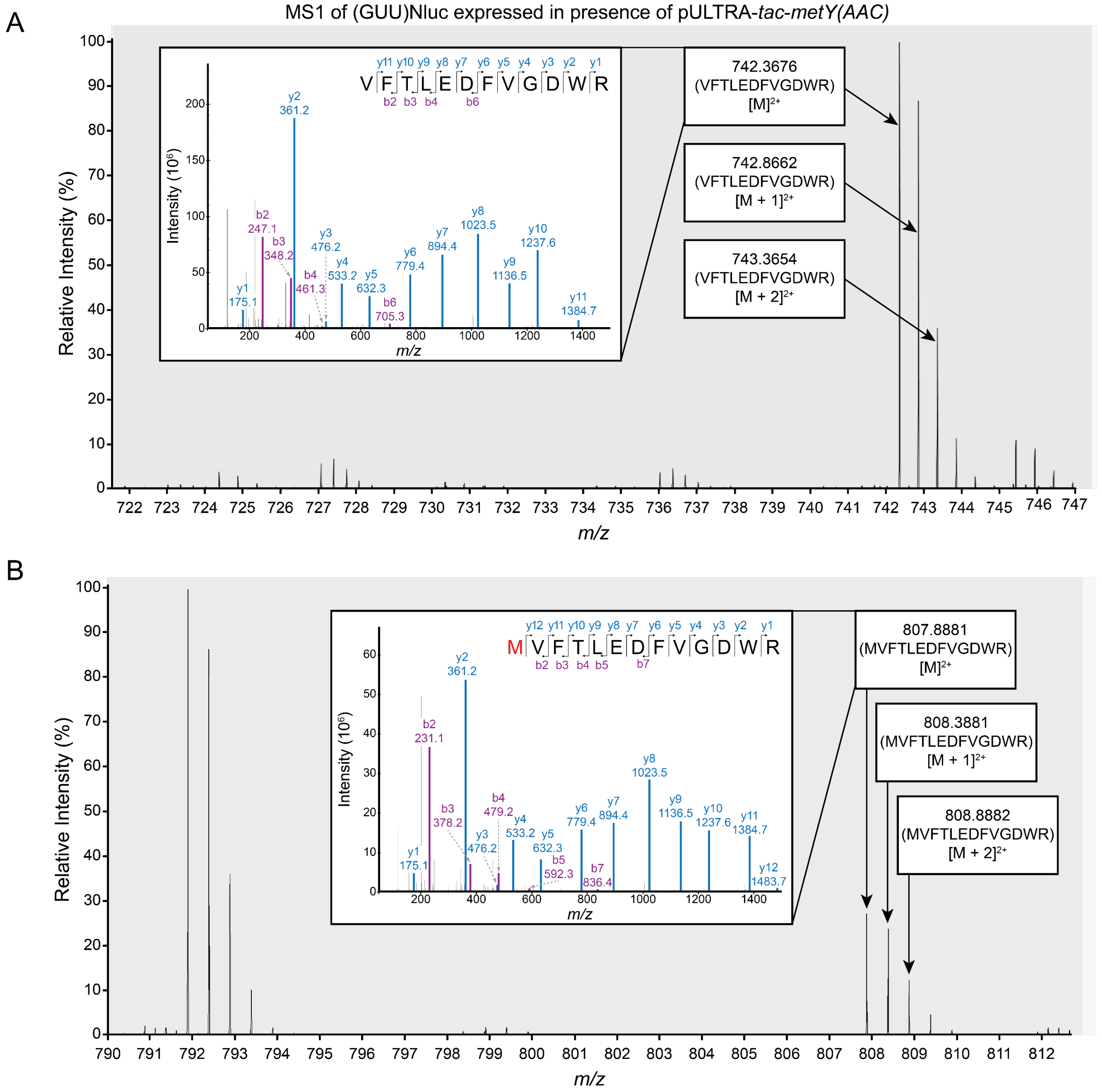


**Figure S7. LC-MS/MS detection of N-terminal peptides of (GUU)Nluc reporter protein purified from *E. coli* BL21(DE3)pLysS (pULTRA-*tac-metY*(AAC); pET20b-*T7*-(*GUU)Nluc6HIS*).** (A) MS1 scan during retention time range of 45-46 minutes. Labelled monoisotopic peak (742.3676 *m/z*) consistent with the N-terminal peptide with residue at N-terminus removed (VFTLEDFVGDWR) and its respective isotopes are shown. The inset shows MS2 scan with y$^-$ and b$^-$ series product ion signals used to confirm amino acid sequence of N-terminal peptide. Schematic of N-terminal peptide (top right of insets) shows peptide fragmentation patterns occurring to produce each of the MS2 ions. (B) MS1 scan during retention time range of 46-7 minutes showing monoisotopic peak (807.8881 *m/z*) consistent with the N-terminal peptide bearing methionine (MVFTLEDFVGDWR) at the N-terminus. MS2 product ions and peptide schematic are shown in inset.

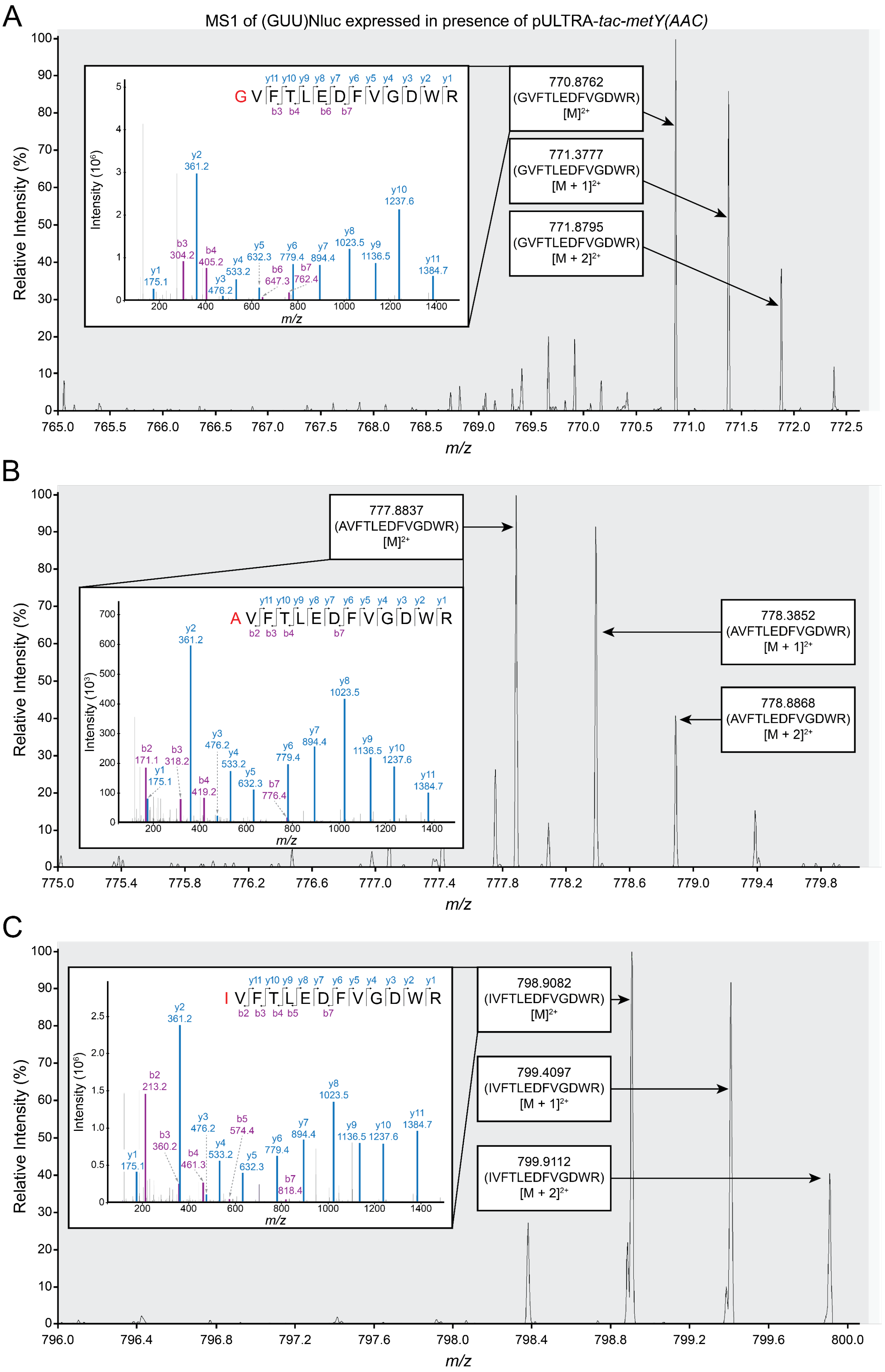
A
MS1 of (GUU)Nluc expressed in presence of pULTRA-tac-metY(AAC)
Relative Intensity (%)
Intensity (10⁶)
G V F T L E D F V G D W R
y11 y10 y9 y8 y7 y6 y5 y4 y3 y2 y1
b3 b4 b6 b7
y1 175.1
b3 304.2
y2 361.2
b4 405.2
y3 476.2
y4 533.2
y5 632.3
b6 647.3
y6 779.4
b7 762.4
y7 894.4
y8 1023.5
y9 1136.5
y10 1237.6
y11 1384.7
m/z
770.8762 (GVFTLEDFVGDWR) [M]2+
771.3777 (GVFTLEDFVGDWR) [M + 1]2+
771.8795 (GVFTLEDFVGDWR) [M + 2]2+
m/z
B
Relative Intensity (%)
777.8837 (AVFTLEDFVGDWR) [M]2+
Intensity (10³)
A V F T L E D F V G D W R
y11 y10 y9 y8 y7 y6 y5 y4 y3 y2 y1
b2 b3 b4 b7
b2 171.1
y1 175.1
b3 318.2
y2 361.2
b4 419.2
y3 476.2
y4 533.2
y5 632.3
b7 776.4
y6 779.4
y7 894.4
y8 1023.5
y9 1136.5
y10 1237.6
y11 1384.7
m/z
778.3852 (AVFTLEDFVGDWR) [M + 1]2+
778.8868 (AVFTLEDFVGDWR) [M + 2]2+
m/z
C
Relative Intensity (%)
Intensity (10⁶)
I V F T L E D F V G D W R
y11 y10 y9 y8 y7 y6 y5 y4 y3 y2 y1
b2 b3 b4 b5 b7
y1 175.1
b2 213.2
b3 360.2
y2 361.2
b4 461.3
y3 476.2
y4 533.2
b5 574.4
y5 632.3
y6 779.4
b7 818.4
y7 894.4
y8 1023.5
y9 1136.5
y10 1237.6
y11 1384.7
m/z
798.9082 (IVFTLEDFVGDWR) [M]2+
799.4097 (IVFTLEDFVGDWR) [M + 1]2+
799.9112 (IVFTLEDFVGDWR) [M + 2]2+
m/z

**Figure S8. LC-MS/MS detection of low abundance N-terminal peptides from (AUG)Nluc reporter protein bearing glycine, alanine, or isoleucine at the N-terminus purified from *E. coli* BL21(DE3)pLysS (pULTRA-*tac-Empty*; pET20b-*T7-(AUG)Nluc6HIS*).** (A) MS1 scan during retention time range of 45-45.5 minutes. Labelled monoisotopic peak (770.8762*m/z*) consistent with an N-terminal peptide bearing glycine (GVFTLEDFVGDWR) and its respective isotopes are shown. he inset shows MS2 scan with $y^-$ and $b^-$ series product ion signals used to confirm amino acid sequence of N-terminal peptide. Schematic of N-terminal peptide (top right of insets) shows peptide fragmentation patterns occurring to produce each of the MS2 ions. (B) MS1 scan during retention time range of 45.2-45.5 minutes showing monoisotopic peak (778.3852 *m/z*) consistent with the N-terminal peptide bearing alanine (AVFTLEDFVGDWR) at the N-terminus. MS2 product ions and peptide schematic are shown in inset. (C) MS1 scan during retention time 46.4-46.9 minutes showing monoisotopic peak (798.9082 *m/z*) consistent with the N-terminal peptide bearing isoleucine (IVFTLEDFVGDWR) at the N-terminus. MS2 product ions and peptides schematic are again shown in the inset.

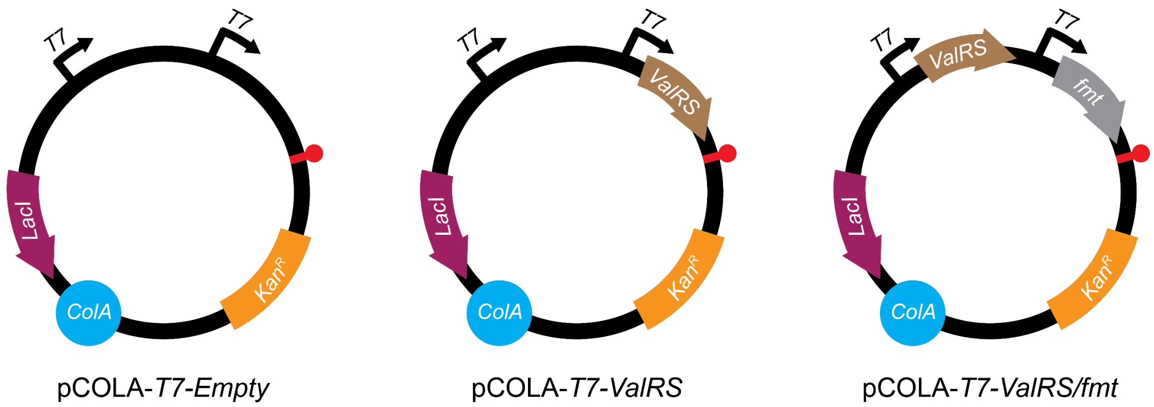


**Figure S9. Schematic of pCOLA plasmids designed to overexpress ValRS or ValRS/FMT.** The pCOLA plasmids contain the ColA origin of replication, Kanamycin resistance gene, two T7 promoter sites, a T7 terminator, and either *ValRS,* or *ValRS and fmt* genes as indicated above.

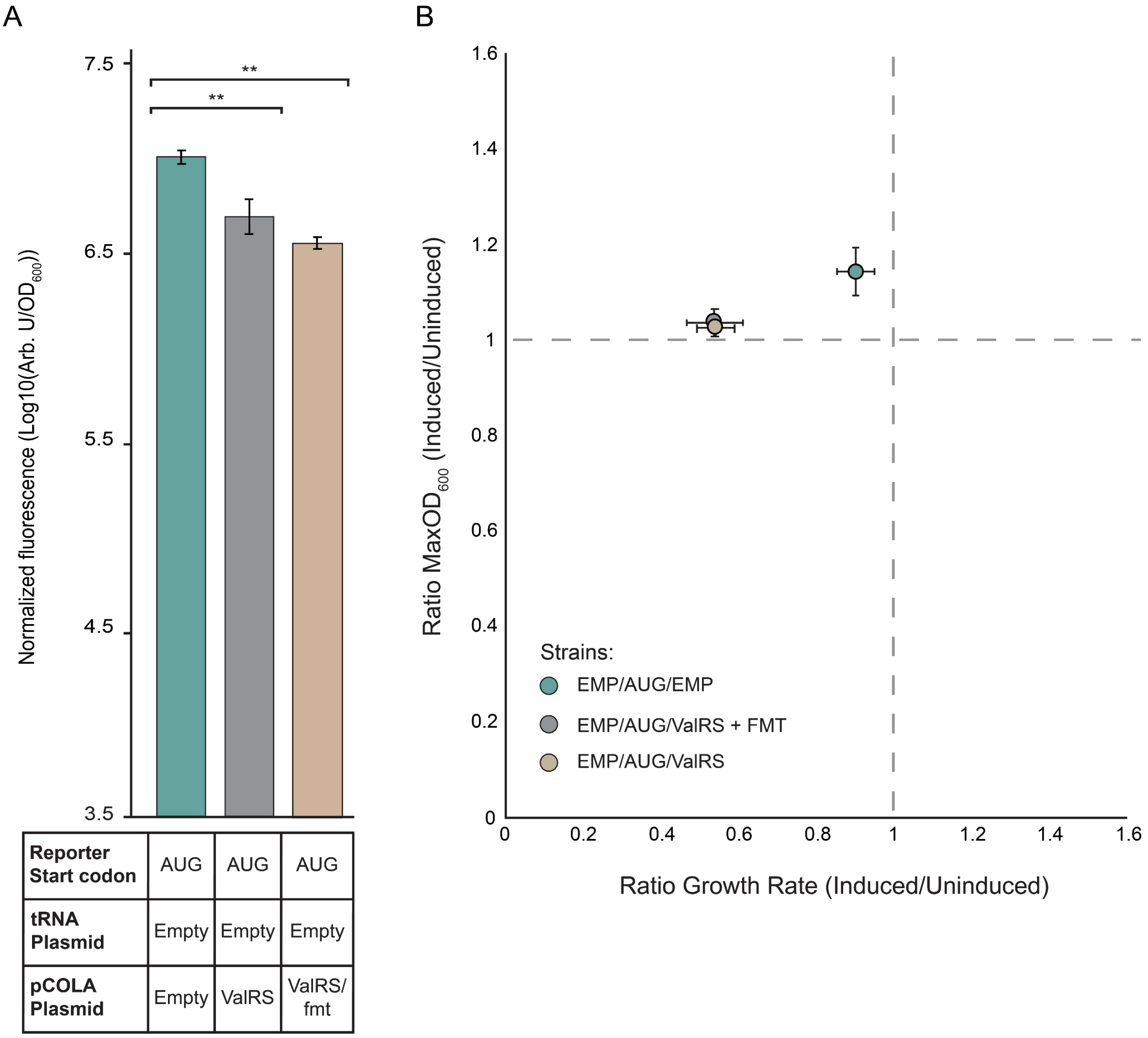


**Figure S10. Reduction in native (AUG)sfGFP expression and strain fitness during ValRS or ValRS + FMT overexpression.** (A) Normalized bulk cell fluorescence (Arb. U./$OD_{600}$) of *E. coli* BL21(DE3)pLysS cells carrying pCOLA-T7-*ValRS* or pCOLA-T7-*ValRS/fmt*. (B) Ratio of $MaxOD_{600}$ and growth rate of *E. coli* BL21(DE3)pLysS cells carrying pULTRA-empty and either pCOLA-T7-*ValRS* or pCOLA-T7-*ValRS/fmt* plasmids.

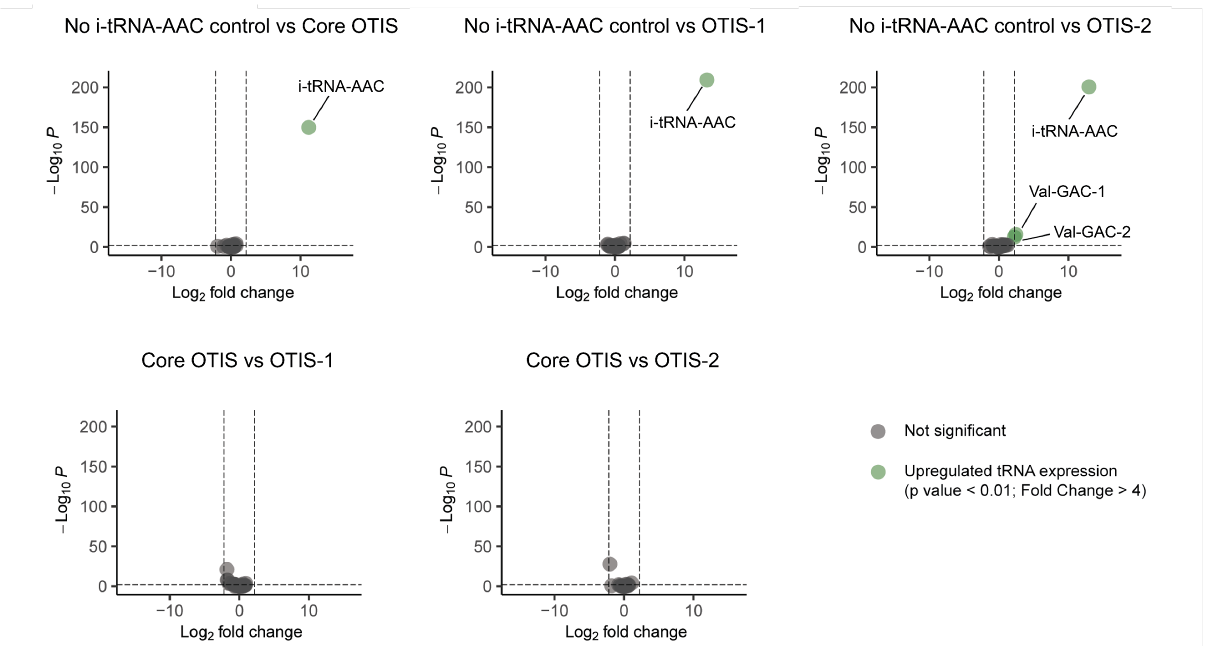


**Figure S11. Differential expression analysis of tRNA measured from *E. coli* BL21(DE3)pLysS cells expressing i-tRNA-AAC with either ValRS, or both ValRS and FMT, compared to cells harbouring an empty pCOLA plasmid.** Vertical gray dashed lines represent fold change threshold: $\log_2$ fold change ≥ 2 or ≤ −2 equivalent to fold change ≥ 4 or ≤ 4, while the horizontal gray dashed line represents significance threshold: $-\log_{10}$ p-value ≥ 2.3 equivalent to p-value ≤ 0.01. Green points represent significantly up-regulated tRNA expression, and gray points represent tRNAs that did not reach the significance threshold (p-value ≥ 0.01).

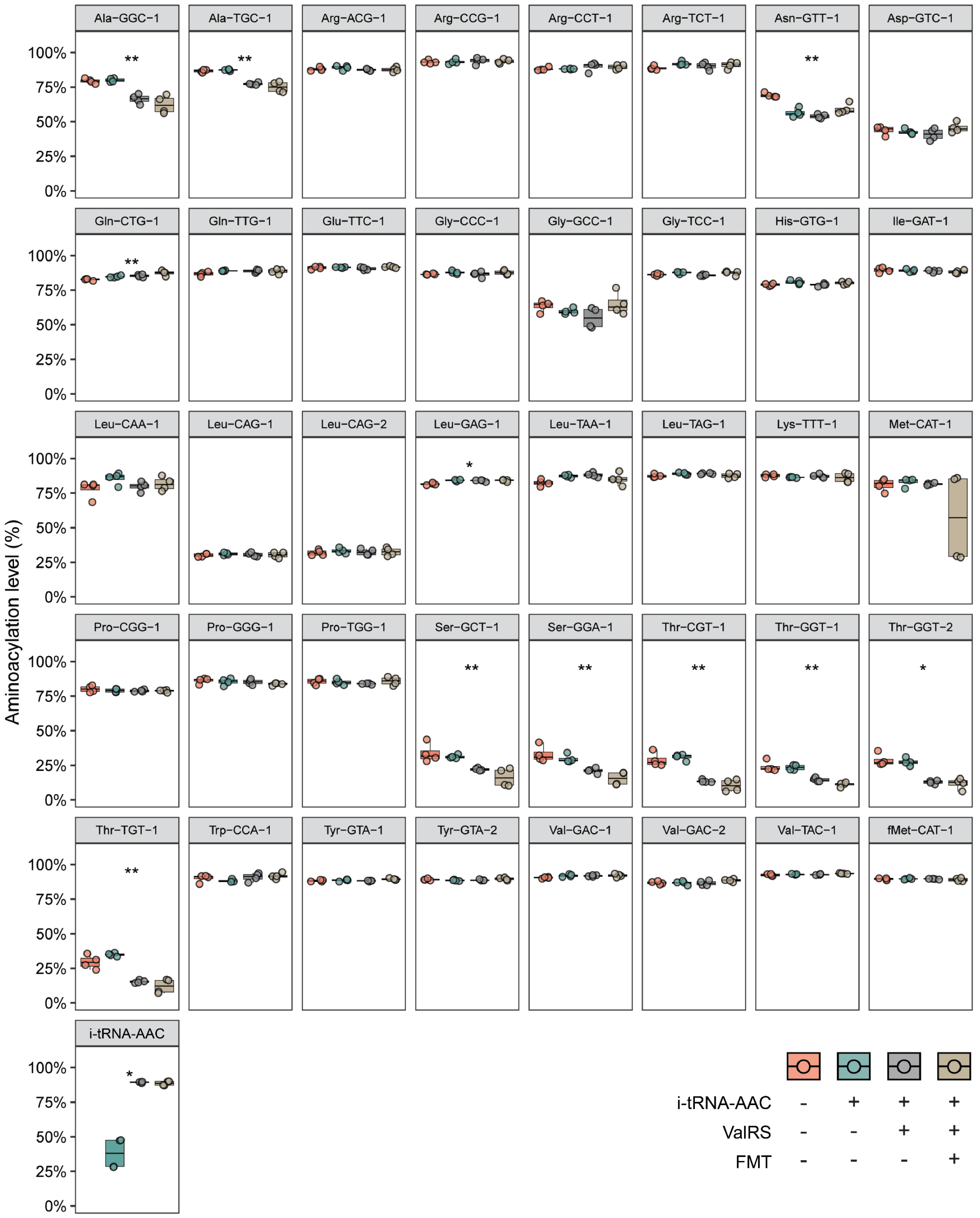


**Figure S12. Aminoacylation level of all tRNA isodecoders.** Measured from *E. coli* BL21(DE3)pLysS cells expressing i-tRNA-AAC with either ValRS, or both ValRS and FMT, compared to cell harbouring an empty pCOLA plasmid. Kruskal Wallis H-Test *p* values, **** < 0.0001, *** < 0.001, ** < 0.01, and * < 0.05.

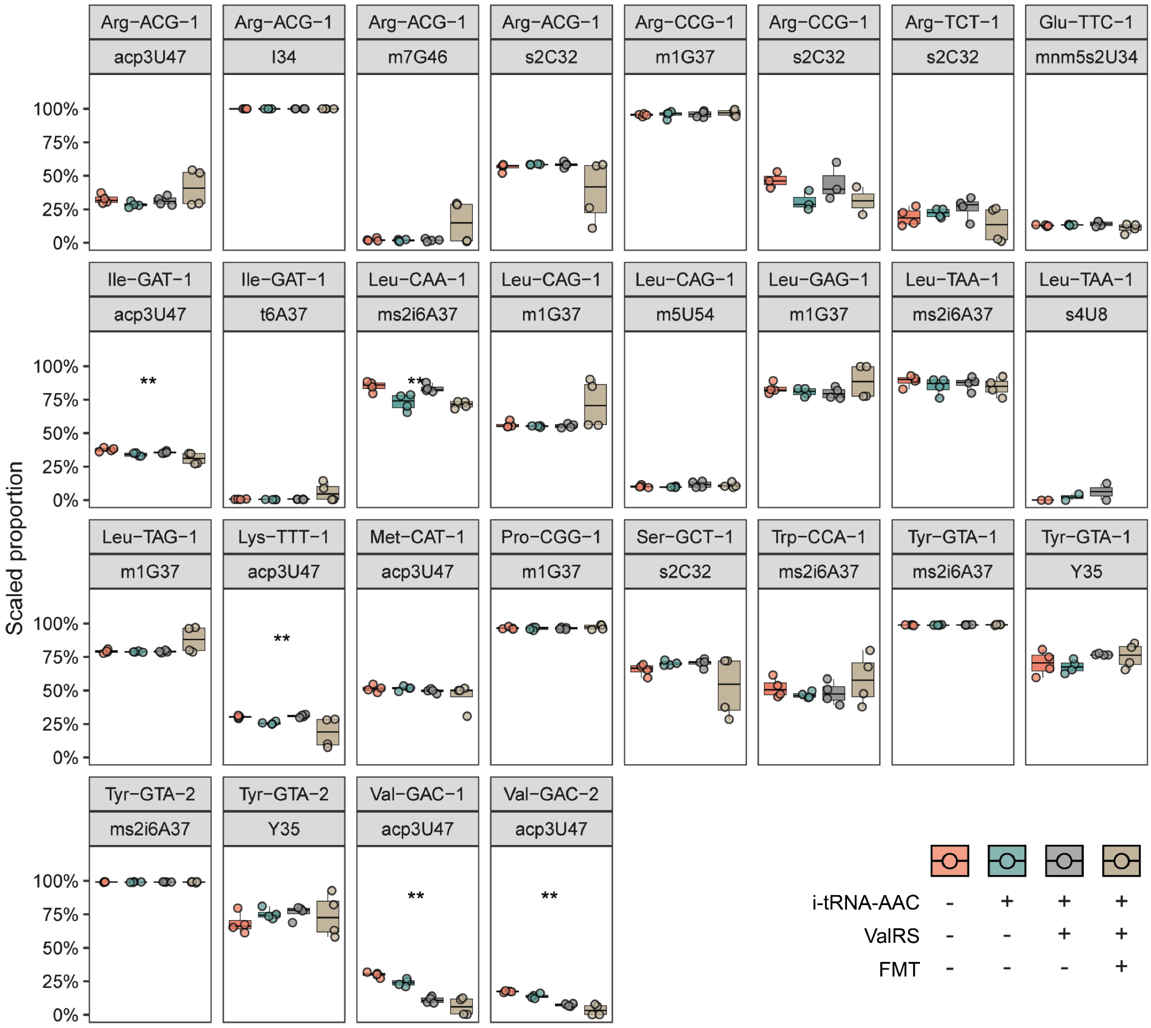


**Figure S13. Expression of ValRS or ValRS/FMT influence tRNA modifications that induce reverse transcription (RT) mismatches.** (A) Scaled proportion of RT misincorporation (%) at canonical tRNA positions displaying > 12.3% mismatch rate. Measured from *E. coli* BL21(DE3)pLysS cells expressing itRNA(AAC) with either ValRS, or both ValRS and FMT, compared to cell harbouring an empty pCOLA plasmid. Kruskal Wallis H-Test *p* values, **** < 0.0001, *** < 0.001, ** < 0.01, and * < 0.05.

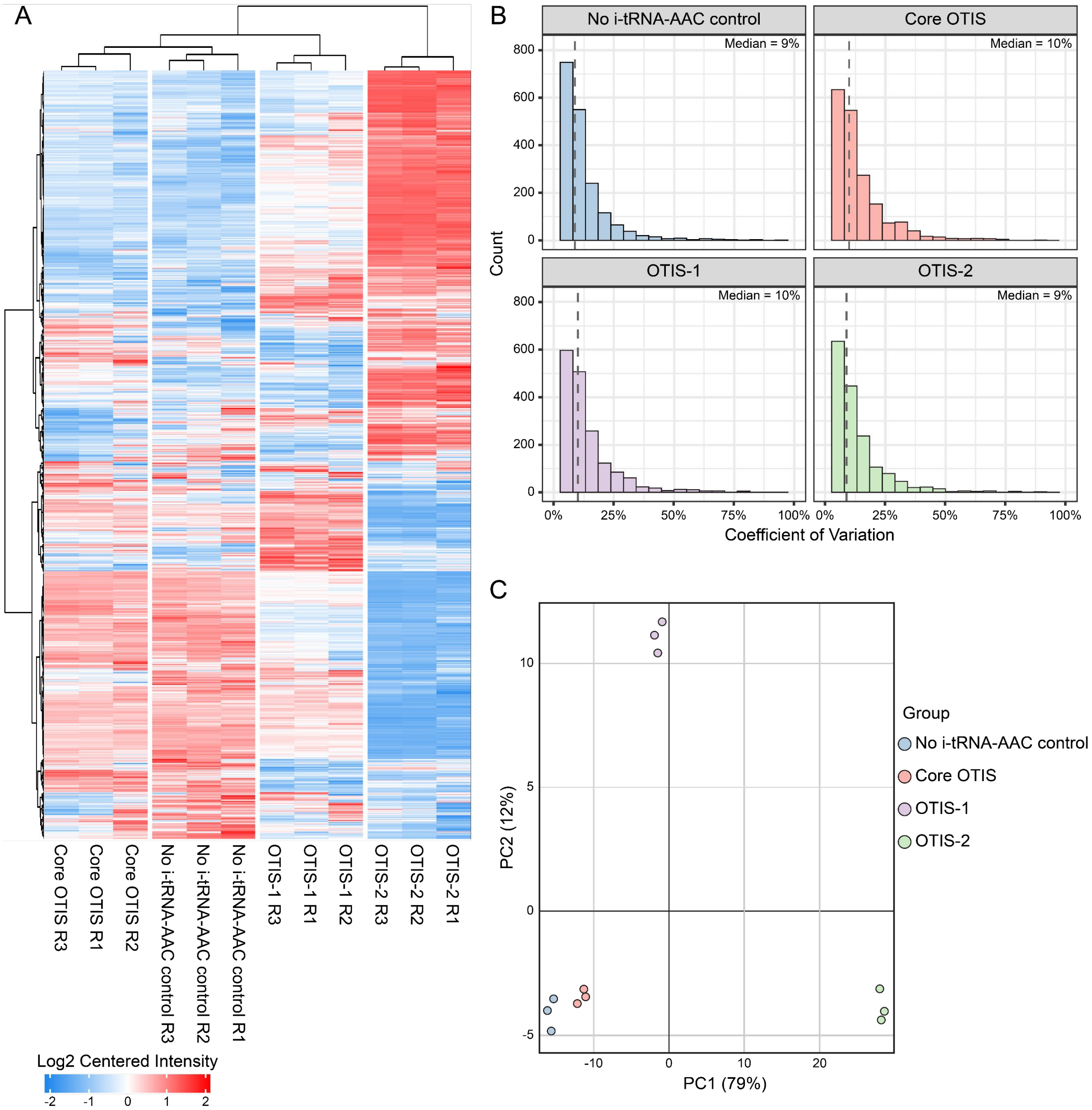


**Figure S14. DIA proteomic analysis of the effects of OTISs**. (A) The hierarchical heatmap of the differentially expressed proteins identified in the DIA-MS based quantitation between the no i-tRNA control, Core OTIS, OTIS-ValRS, and OTIS-ValRS/FMT overexpressed strains. (B) Bar graph showing the Coefficient of variation in each of the sample time. (C) Principal component analysis of each the replicates of each strain category.

### SUPPORTING TABLES

**Table S1 ICM-Pro ligand docking scores for peptides fMAS, fVAS and fAAS**

| *PEPTIDE* | EDOCK | EGB | EGE | EGS | EGV |
|---|---|---|---|---|---|
| ***fMAS*** | -32.27 | -5.96 | -12.04 | -1.57 | -61.88 |
| ***fVAS*** | -30.47 | -7.18 | -13.46 | -1.45 | -57.49 |
| ***fAAS*** | -27.73 | -7.17 | -13.08 | -0.74 | -51.01 |

fMAS, formylated-methionine-alanine-serine peptide; fVAS, formylated-valine; fAAS, formylated-alanine. Edoc, docking score where lower values represent better binding. It is a composite score based on hydrogen bond grid energy (Egb), electrostatic grid potential (Ege), hydrophobic potential (Egs) and van der Waals interaction (Egv).

**Table S2. ICM-Pro peptide docking scores fMAS.**

| *Peptide* | Edock | Egb | Ege | Egs | Egv |
|---|---|---|---|---|---|
| *fMAS_1* | -32.27 | -5.96 | -12.04 | -1.57 | -61.88 |
| *fMAS_2* | -29.59 | -11.80 | -16.35 | -1.23 | -53.45 |
| *fMAS_3* | -25.87 | -6.70 | -10.40 | -1.56 | -58.50 |
| *fMAS_4* | -24.30 | -10.39 | -11.58 | -1.40 | -53.61 |
| *fMAS_5* | -24.10 | -8.28 | -12.37 | -2.29 | -52.42 |
| *fMAS_6* | -22.37 | -7.99 | -5.18 | -0.58 | -62.31 |
| *fMAS_7* | -21.70 | -8.04 | -7.70 | -0.54 | -59.63 |
| *fMAS_8* | -21.27 | -14.27 | -13.02 | -1.53 | -43.71 |
| *fMAS_9* | -21.23 | -7.02 | -12.91 | -1.81 | -52.36 |
| *fMAS_10* | -21.19 | -14.31 | -14.61 | -1.35 | -43.70 |

**Table S3. ICM-Pro peptide docking scores fVAS.**

| *Peptide* | Edock | Egb | Ege | Egs | Egv |
|---|---|---|---|---|---|
| *fVAS_1* | -30.47 | -7.18 | -13.46 | -1.45 | -57.49 |
| *fVAS_2* | -28.39 | -9.31 | -7.45 | -0.22 | -57.17 |
| *fVAS_3* | -26.58 | -10.57 | -12.31 | -1.53 | -48.77 |
| *fVAS_4* | -25.85 | -12.77 | -16.82 | -1.42 | -46.69 |
| *fVAS_5* | -22.88 | -13.63 | -16.08 | -1.82 | -40.81 |
| *fVAS_6* | -22.82 | -13.18 | -17.79 | -1.56 | -38.50 |
| *fVAS_7* | -20.78 | -13.57 | -17.80 | -1.41 | -31.60 |
| *fVAS_8* | -20.39 | -7.85 | -9.13 | 1.03 | -52.82 |
| *fVAS_9* | -20.20 | -10.46 | -5.61 | -0.39 | -51.24 |
| *fVAS_10* | -20.14 | -7.59 | -3.89 | 0.42 | -58.84 |

**Table S4. tRNA positions from mim-tRNAseq dataset with more than 10% reverse-transcription misincorporation.**

| tRNA | Canonical Position | Base Identity | Modification | Empty | Core OTIS | OTIS-1 | OTIS-2 |
|---|---|---|---|---|---|---|---|
| Arg-ACG-1 | 32 | C | 2-thiocytidine | 56.29 ± 2.9% | 58.46 ± 0.27% | 58.33 ± 2.12% | 38.19 ± 23.55% |
| Arg-ACG-1 | 34 | A | inosine | 100 ± 0% | 100 ± 0% | 100 ± 0% | 100 ± 0% |
| Arg-ACG-1 | 46 | G | 7-methylguanosine | 2.21 ± 1.06% | 1.76 ± 0.72% | 1.93 ± 0.96% | 15.1 ± 15.96% |
| Arg-ACG-1 | 47 | U | 3-(3-amino-3-carboxypropyl)uridine | 32.53 ± 3.5% | 28.51 ± 2.01% | 31.4 ± 3.37% | 41.1 ± 14% |
| Arg-CCG-1 | 32 | C | 2-thiocytidine | 46.67 ± 6.03% | 30.9 ± 7.35% | 44.44 ± 13.88% | 31.36 ± 14.58% |
| Arg-CCG-1 | 37 | G | 1-methylguanosine | 95.41 ± 0.99% | 95.68 ± 2.62% | 95.92 ± 2.46% | 96.93 ± 2.45% |
| Arg-TCT-1 | 32 | C | 2-thiocytidine | 19.36 ± 6.91% | 22.11 ± 3.33% | 26.06 ± 8.53% | 13.44 ± 13.34% |
| Glu-TTC-1 | 34 | U | 5-methylaminomethyl-2-thiouridine | 12.91 ± 0.56% | 13.34 ± 0.36% | 14.11 ± 1.88% | 10.91 ± 3.45% |
| Ile-GAT-1 | 37 | A | N6-threonylcarbamoyladenosine | 0.72 ± 0.16% | 0.53 ± 0.09% | 0.69 ± 0.21% | 6.04 ± 6.81% |
| Ile-GAT-1 | 47 | U | 3-(3-amino-3-carboxypropyl)uridine | 37.67 ± 1.46% | 34.03 ± 1.31% | 35.67 ± 0.8% | 31.05 ± 4.32% |
| Leu-CAA-1 | 37 | A | 2-methylthio-N6-isopentenyladenosine | 84.97 ± 3.9% | 73.01 ± 6.35% | 83.41 ± 3.02% | 71.31 ± 2.7% |
| Leu-CAG-1 | 37 | G | 1-methylguanosine | 56.19 ± 2.33% | 55.1 ± 0.62% | 55.4 ± 1.57% | 71.87 ± 18.3% |
| Leu-CAG-1 | 54 | U | 5-methyluridine | 10.26 ± 1.02% | 9.87 ± 0.49% | 11.68 ± 2.41% | 11 ± 1.88% |
| Leu-GAG-1 | 37 | G | 1-methylguanosine | 83.29 ± 4.05% | 80.65 ± 3.12% | 79.86 ± 4.23% | 88.58 ± 12.78% |
| Leu-TAA-1 | 37 | A | 2-methylthio-N6-isopentenyladenosine | 88.86 ± 4.35% | 85.04 ± 6.45% | 86.97 ± 5.05% | 84.52 ± 6.98% |
| Leu-TAA-1 | 8 | U | 4-thiouridine | 0 ± 0% | 2.27 ± 3.21% | 6.25 ± 8.84% | NA |
| Leu-TAG-1 | 37 | G | 1-methylguanosine | 79.17 ± 1.37% | 78.92 ± 0.47% | 79.04 ± 0.89% | 88.08 ± 9.99% |
| Lys-TTT-1 | 47 | U | 3-(3-amino-3-carboxypropyl)uridine | 30.26 ± 0.93% | 25.65 ± 0.98% | 30.98 ± 1% | 18.6 ± 11.31% |

| | | | | | | | |
|---|---|---|---|---|---|---|---|
| Met-CAT-1 | 47 | U | 3-(3-amino-3-carboxypropyl)uridine | 51.36 ± 2.6% | 51.58 ± 1.73% | 49.68 ± 1.61% | 45.62 ± 9.94% |
| Pro-CGG-1 | 37 | G | 1-methylguanosine | 96.58 ± 0.88% | 96.26 ± 1.16% | 96.41 ± 0.81% | 97.3 ± 1.81% |
| Ser-GCT-1 | 32 | C | 2-thiocytidine | 65.54 ± 4.43% | 70.26 ± 1.76% | 70.41 ± 3.3% | 52.54 ± 22.87% |
| Trp-CCA-1 | 37 | A | 2-methylthio-N6-isopentenyladenosine | 51.96 ± 7.3% | 46.61 ± 2.29% | 48.17 ± 8.46% | 58.27 ± 19.11% |
| Tyr-GTA-1 | 35 | U | pseudouridine | 70.36 ± 9.29% | 67.82 ± 4.83% | 76.81 ± 0.65% | 75.89 ± 9.41% |
| Tyr-GTA-1 | 37 | A | 2-methylthio-N6-isopentenyladenosine | 98.78 ± 0.23% | 98.79 ± 0.18% | 98.91 ± 0.12% | 98.96 ± 0.32% |
| Tyr-GTA-2 | 35 | U | pseudouridine | 68.37 ± 7.95% | 75.32 ± 4.09% | 76.39 ± 5.1% | 74 ± 16.18% |
| Tyr-GTA-2 | 37 | A | 2-methylthio-N6-isopentenyladenosine | 99.09 ± 0.2% | 99.04 ± 0.21% | 99.1 ± 0.07% | 99 ± 0.36% |
| Val-GAC-1 | 47 | U | 3-(3-amino-3-carboxypropyl)uridine | 29.98 ± 1.97% | 23.93 ± 2.61% | 11.03 ± 2.43% | 6.23 ± 6.71% |
| Val-GAC-2 | 47 | U | 3-(3-amino-3-carboxypropyl)uridine | 17.38 ± 0.62% | 13.9 ± 1.67% | 7.35 ± 1.02% | 3.65 ± 4.07% |

**Table S5. Gene sequences used in this study.**

| Gene | Sequence (5’-3’) |
|---|---|
| *sfGFP* | ATGCGTAAAGGCGAAGAGCTGTTCACTGGTGTCGTCCCTATTCTGGTGGAACTG<br>GATGGTGATGTCAACGGTCATAAGTTTTCCGTGCGTGGCGAGGGTGAAGGTGAC<br>GCAACTAATGGTAAACTGACGCTGAAGTTCATCTGTACTACTGGTAAACTGCCGG<br>TACCTTGGCCGACTCTGGTAACGACGCTGACTTATGGTGTTCAGTGCTTTGCTCG<br>TTATCCGGACCATATGAAGCAGCATGACTTCTTCAAGTCCGCCATGCCGGAAGGC<br>TATGTGCAGGAACGCACGATTTCCTTTAAGGATGACGGCACGTACAAAACGCGT<br>GCGGAAGTGAAATTTGAAGGCGATACCCTGGTAAACCGCATTGAGCTGAAAGGC<br>ATTGACTTTAAAGAAGACGGCAATATCCTGGGCCATAAGCTGGAATACAATTTTA<br>ACAGCCACAATGTTTACATCACCGCCGATAAACAAAAAAATGGCATTAAAGCGA<br>ATTTTAAAATTCGCCACAACGTGGAGGATGGCAGCGTGCAGCTGGCTGATCACT<br>ACCAGCAAAACACTCCAATCGGTGATGGTCCTGTTCTGCTGCCAGACAATCACTA<br>TCTGAGCACGCAAAGCGTTCTGTCTAAAGATCCGAACGAGAAACGCGATCATAT<br>GGTTCTGCTGGAGTTCGTAACCGCAGCGGGCATCACGCATGGTATGGATGAACT<br>GTACAAATGATGA |
| *Nluc6xHIS* | ATGGTCTTCACACTCGAAGATTTCGTTGGGGACTGGCGACAGACAGCCGGCTAC<br>AACCTGGACCAAGTCCTTGAACAGGGAGGTGTGTCCAGTTTGTTTCAGAATCTCG<br>GGGTGTCCGTAACTCCGATCCAAAGGATTGTCCTGAGCGGTGAAAATGGGCTGA<br>AGATCGACATCCATGTCATCATCCCGTATGAAGGTCTGAGCGGCGACCAAATGG<br>GCCAGATCGAAAAAATTTTTAAGGTGGTGTACCCTGTGGATGATCATCACTTTAA<br>GGTGATCCTGCACTATGGCACACTGGTAATCGACGGGGTTACGCCGAACATGAT<br>CGACTATTTCGGACGGCCGTATGAAGGCATCGCCGTGTTCGACGGCAAAAAGAT<br>CACTGTAACAGGGACCCTGTGGAACGGCAACAAAATTATCGACGAGCGCCTGAT<br>CAACCCCGACGGCTCCCTGCTGTTCCGAGTAACCATCAACGGAGTGACCGGCTG<br>GCGGCTGTGCGAACGCATTCTGGCGCACCACCACCACCACCACTAATAA |
| *metY(AAC)* | CGCGGGGTGGAGCAGCCTGGTAGCTCGTCGGGCTAACAACCCGAAGATCGTCG<br>GTTCAAATCCGGCCCCCGCAACCA |
| *fmt* | ATGTCAGAATCACTACGTATTATTTTTGCGGGTACACCTGACTTTGCAGCGCGTC<br>ATCTCGACGCGCTGTTGTCTTCTGGTCATAACGTCGTTGGCGTGTTCACCCAGCC<br>AGACCGACCGGCAGGACGCGGTAAAAAACTGATGCCCAGCCCGGTTAAAGTTCT<br>GGCTGAGGAAAAAGGTCTGCCCGTTTTTCAACCTGTTTCCCTGCGTCCACAAGAA<br>AACCAGCAACTGGTCGCCGAACTGCAGGCTGATGTTATGGTCGTCGTCGCCTAT<br>GGTTTAATTCTGCCGAAAGCAGTGCTGGAGATGCCGCGTCTTGGCTGTATCAAC<br>GTTCATGGTTCACTGCTGCCACGCTGGCGCGGTGCTGCACCAATCCAACGCTCAC<br>TATGGGCGGGTGATGCAGAAACTGGTGTGACCATTATGCAAATGGATGTCGGTT<br>TAGACACCGGTGATATGCTCTATAAGCTCTCCTGCCCGATTACTGCAGAAGATAC<br>CAGTGGTACGCTGTACGACAAGCTGGCAGAGCTTGGCCCACAAGGGCTTATCAC<br>CACGTTGAAACAACTGGCAGACGGCACGGCGAAACCAGAAGTTCAGGACGAAA<br>CTCTTGTCACTTACGCCGAGAAGTTGAGTAAAGAAGAAGCGCGTATTGACTGGT<br>CACTTTCGGCAGCACAGCTTGAACGCTGCATTCGCGCTTTCAATCCATGGCCAAT<br>GAGCTGGCTGGAAATTGAAGGACAGCCGGTTAAAGTCTGGAAAGCATCGGTCAT<br>TGATACGGCAACCAACGCTGCACCAGGAACGATCCTTGAAGCCAACAAACAAGG<br>CATTCAGGTTGCGACTGGTGATGGCATCCTGAACCTGCTCTCGTTACAACCTGCG |

| | |
|---|---|
| | GGTAAGAAAGCGATGAGCGCGCAAGACCTCCTGAACTCTCGTCGGGAATGGTTT<br>GTTCCGGGCAACCGTCTGGTCTGA |
| *ValS* | ATGGAAAAGACATATAACCCACAAGATATCGAACAGCCGCTTTACGAGCACTGG<br>GAAAAGCAGGGCTACTTTAAGCCTAATGGCGATGAAAGCCAGGAAAGTTTCTGC<br>ATCATGATCCCGCCGCCGAACGTCACCGGCAGTTTGCATATGGGTCACGCCTTCC<br>AGCAAACCATCATGGATACCATGATCCGCTATCAGCGCATGCAGGGCAAAAACA<br>CCCTGTGGCAGGTCGGTACTGACCACGCCGGGATCGCTACCCAGATGGTCGTTG<br>AGCGCAAGATTGCCGCAGAAGAAGGTAAAACCCGTCACGACTACGGCCGCGAA<br>GCTTTCATCGACAAAATCTGGGAATGGAAAGCGGAATCTGGCGGCACCATTACC<br>CGTCAGATGCGCCGTCTCGGCAACTCCGTCGACTGGGAGCGTGAACGCTTCACC<br>ATGGACGAAGGCCTGTCCAATGCGGTGAAAGAAGTTTTCGTTCGTCTGTATAAA<br>GAAGACCTGATTTACCGTGGCAAACGCCTGGTAAACTGGGATCCGAAACTGCGC<br>ACCGCTATCTCTGACCTGGAAGTCGAAAACCGCGAATCGAAAGGTTCGATGTGG<br>CACATCCGCTATCCGCTGGCTGACGGTGCGAAAACCGCAGACGGTAAAGATTAT<br>CTGGTGGTCGCGACTACCCGTCCAGAAACCCTGCTGGGCGATACTGGCGTAGCC<br>GTTAACCCGGAAGATCCGCGTTACAAAGATCTGATTGGCAAATATGTCATTCTGC<br>CGCTGGTTAACCGTCGTATTCCGATCGTTGGCGACGAACACGCCGACATGGAAA<br>AAGGCACCGGCTGCGTGAAAATCACTCCGGCGCACGACTTTAACGACTATGAAG<br>TGGGTAAACGTCACGCCCTGCCGATGATCAACATCCTGACCTTTGACGGCGATAT<br>CCGTGAAAGCGCCCAGGTGTTCGATACCAAAGGTAACGAATCTGACGTTTATTCC<br>AGCGAAATCCCTGCAGAGTTCCAGAAACTGGAGCGTTTTGCTGCACGTAAAGCA<br>GTCGTTGCCGCAGTTGACGCGCTTGGCCTGCTGGAAGAAATTAAACCGCACGAC<br>CTGACCGTTCCTTACGGCGACCGTGGCGGCGTAGTTATCGAACCAATGCTGACC<br>GACCAGTGGTACGTGCGTGCCGATGTCCTGGCGAAACCGGCGGTTGAAGCGGTT<br>GAGAACGGCGACATTCAGTTCGTACCGAAGCAGTACGAAAACATGTACTTCTCCT<br>GGATGCGCGATATTCAGGACTGGTGTATCTCTCGTCAGTTGTGGTGGGGTCACC<br>GTATCCCGGCATGGTATGACGAAGCGGGTAACGTTTATGTTGGCCGCAACGAAG<br>ACGAAGTGCGTAAAGAAAATAACCTCGGTGCTGATGTTGTCCTGCGTCAGGACG<br>AAGACGTTCTCGATACCTGGTTCTCTTCTGCGCTGTGGACCTTCTCTACCCTTGGC<br>TGGCCGGAAAATACCGACGCCCTGCGTCAGTTCCACCCAACCAGCGTGATGGTA<br>TCTGGTTTCGACATCATTTTCTTCTGGATTGCCCGCATGATCATGATGACCATGCA<br>CTTCATCAAAGATGAAAATGGCAAACCGCAGGTGCCGTTCCACACCGTTTACATG<br>ACCGGCCTGATTCGTGATGACGAAGGCCAGAAGATGTCCAAATCCAAGGGTAAC<br>GTTATCGACCCACTGGATATGGTTGACGGTATTTCGCTGCCAGAACTGCTGGAAA<br>AACGTACCGGCAATATGATGCAGCCGCAGCTGGCGGACAAAATCCGTAAGCGCA<br>CCGAGAAGCAGTTCCCGAACGGTATTGAGCCGCACGGTACTGACGCGCTGCGCT<br>TCACCCTGGCGGCGCTGGCGTCTACCGGTCGTGACATCAACTGGGATATGAAGC<br>GTCTGGAAGGTTACCGTAACTTCTGTAACAAGCTGTGGAACGCCAGCCGCTTTGT<br>GCTGATGAACACAGAAGGTCAGGATTGCGGCTTCAACGGCGGCGAAATGACGC<br>TGTCGCTGGCGGACCGCTGGATTCTGGCGGAGTTCAACCAGACCATCAAAGCGT<br>ACCGCGAAGCGCTGGACAGCTTCCGCTTCGATATCGCCGCAGGCATTCTGTATGA<br>GTTCACCTGGAACCAGTTCTGTGACTGGTATCTCGAGCTGACCAAGCCGGTAATG<br>AACGGTGGCACCGAAGCAGAACTGCGCGGTACTCGCCATACGCTGGTGACTGTA<br>CTGGAAGGTCTGCTGCGCCTCGCGCATCCGATCATTCCGTTCATCACCGAAACCA<br>TCTGGCAGCGTGTGAAAGTACTTTGCGGTATCACTGCCGACACCATCATGCTGCA<br>GCCGTTCCCGCAGTACGATGCATCTCAGGTTGATGAAGCCGCACTGGCCGACAC<br>CGAATGGCTGAAACAGGCGATCGTTGCGGTACGTAACATCCGTGCAGAAATGAA |

| | |
|---|---|
| | CATCGCGCCGGGCAAACCGCTGGAGCTGCTGCTGCGTGGTTGCAGCGCGGATGC<br>AGAACGTCGCGTAAATGAAAACCGTGGCTTCCTGCAAACCCTGGCGCGTCTGGA<br>AAGTATCACCGTGCTGCCTGCCGATGACAAAGGTCCGGTTTCCGTTACGAAGATC<br>ATCGACGGTGCAGAGCTGCTGATCCCGATGGCTGGCCTCATCAACAAAGAAGAT<br>GAGCTGGCGCGTCTGGCGAAAGAAGTGGCGAAGATTGAAGGTGAAATCAGCCG<br>TATCGAGAACAAACTGGCGAACGAAGGCTTTGTCGCCCGCGCACCGGAAGCGGT<br>CATCGCGAAAGAGCGTGAGAAGCTGGAAGGCTATGCGGAAGCGAAAGCGAAAC<br>TGATTGAACAGCAGGCTGTTATCGCCGCGCTGTAA |

**Table S6. Oligos used (sequencing, Gibson assembly, colony PCR for all plasmids).**

| Oligo | Sequence (5'-3') |
|---|---|
| *metY* Sequencing and colony PCR Primer Forward | TCTCCCTTATGCGACTCCTG |
| *metY* Sequencing and colony PCR primer Reverse | AGATCCGGCCACGATGAC |
| *sfgfp* Sequencing and colony PCR Primer Forward | CTCGCGTATCGGTGATTCAT |
| *sfgfp* Sequencing and colony PCR Primer Reverse | CGGATAACGAGCAAAGCACT |
| *pULTRA* linearisation primer Forward | CGCAACCACTTTCCCTTAGA |
| *pULTRA* linearisation primer Reverse | TGAAAGCACCTCCTTTGTGA |
| *nluc* Sequencing and colony PCR Primer Forward | CGCGTTTCCAGACTTTACG |
| *nluc* Sequencing and colony PCR Primer Reverse | GCTTAATGCGCCGCTACAG |
| *pCOLA* linearisation primer (Site 1) Forward | TGCTTAAGTCGAACAGAAAGT |
| *pCOLA* linearisation primer (Site 1) Reverse | GGTATATCTCCTTATTAAAGTTAAACAA |
| *pCOLA* linearisation primer (Site 2) Forward | TCGAGTCTGGTAAAGAAACC |
| *pCOLA* linearisation primer (Site 2) Reverse | TGTATATCTCCTTCTTATACTTAACTAAT |
| *fmt* genome amplification primer (Site 2 specific handles) Forward | gtataagaaggagatatacaATGTCAGAATCACTACGTATTATTT |
| *fmt* genome amplification primer (Site 2 specific handles) Reverse | ggtttctttaccagactcgaCTTAGAAGAGTGGACTATCAGACC |
| *ValS* genome amplification primer (Site 2 specific handles) Forward | gtataagaaggagatatacaATGGAAAAGACATATAACCCACAAG |

| | |
|---|---|
| *ValS* genome amplification primer (Site 2 specific handles) Reverse | ggtttctttaccagactcgaTCATCACTGTGTTTTGATTACA |
| *ValS* genome amplification primer (Site 1 specific handles) Forward | ctttaataaggagatataccATGGAAAAGACATATAACCCACAAG |
| *ValS* genome amplification primer (Site 1 specific handles) Reverse | actttctgttcgacttaagcaTCATCACTGTGTTTTGATTACA |

**Supplementary Methods:**

To model the relative binding stability of formylated XAS mutant peptides, we used the $Ni^{2+}$ bound enzyme structure (pdb code: 1bs6) as this is has been shown to be an active deformylase[2] and has been defined bound to partial substrate peptide MAS[3]. Molsoft ICM-Pro software was used to calculate the relative difference in binding free energy ($\Delta\Delta G$) between mutant and wildtype complexes, computed from the sum of van der Waals attraction and repulsion, electrostatic interactions, change in entropy, hydrogen bond formation and solvent interactions using the Internal Coordinate Force Field (ICFF)[4]. A Monte Carlo simulation of side chain repacking in the vicinity of the mutated residue was performed to optimise the lowest energy mutant conformation and binding interaction while maintaining the geometry of the peptide backbone[5-7].

**Supplementary Discussion:**

A more detailed prediction of possible binding poses was performed by docking models of formylated substrates fMAS, fVAS and fAAS into the Ni-bound active site of peptide deformylase (1bs6, chain A). The docking score (Edoc) represents a composite score where the lower the score, the better the binding, and is calculated from a combination of hydrogen bond grid energy (Egb), electrostatic grid potential (Ege), hydrophobic potential (Egs) and van der Waals interaction (Egv) (Tables S2-S3). The best scoring example of each peptide indicates a binding preference in the order fMAS>fVAS>fAAS (Edoc scores) (main text, Table 2). This correlated with Egv and Egs scores, which demonstrated the highest variance between mutant peptides for the best poses, indicating that van der Waals and hydrophobic interactions are likely the basis of substrate selectivity. Docking was performed in triplicate; and the most negative scoring docked peptides show very limited variance in backbone architecture proximal to the active site (main text Fig. 5A), with the fMAS peptide having the carbonyl oxygen of the formyl group coordinated in proximity to the catalytic nucleophile W1 (3.1 Å), the backbone amide hydrogen of L91 (2.3 Å), and the active site Ni atom (3.4 Å) (main text Fig. 5B), consistent with the mechanism of peptide deformylase formyl hydrolysis proposed by Becker et al. [3] and modelled by Wu et al. [8]. The highest scoring docked fVAS peptide shows a similar orientation, with the formyl group carbonyl oxygen more closely associated with W1 (2.8 Å) and Ni (3.0 Å), and slightly further from L91 (2.7 Å) (main text Fig. 5C). Notably, in the crystal structure of unformylated peptide (MAS) bound enzyme, W2 occupies the equivalent site to the formyl oxygen in the fMAS and fVAS top docked poses. No profound differences in the active site localisation of the formyl group are evident, indicating that this binding event would be competent for catalysis. The molecular determinants of specificity are clearer when rendered as a 2-dimensional interaction diagram. A decrease in hydrophobic interactions is evident for fVAS, relative to fMAS (Fig. 5D-E). The progressive decrease in accessible surface area for the side chain at position 1 may explain the apparent decreased catalytic efficiency in hydrolysis of the formyl group of formyl-Valine bearing proteins, evidenced by their detection by mass spectrometry, and the absence of formylated methionine in this analysis.

**Supplementary References**

[1] Chan PP, Lowe TM. GtRNAdb: a database of transfer RNA genes detected in genomic sequence. Nucleic acids research. 2009;37:D93-D7.
[2] Groche D, Becker A, Schlichting I, Kabsch W, Schultz S, Wagner AV. Isolation and Crystallization of Functionally CompetentEscherichia coliPeptide Deformylase Forms Containing either Iron or Nickel in the Active Site. Biochemical and biophysical research communications. 1998;246:342-6.
[3] Becker A, Schlichting I, Kabsch W, Groche D, Schultz S, Wagner AV. Iron center, substrate recognition and mechanism of peptide deformylase. Nature structural biology. 1998;5:1053-8.
[4] Katritch V, Totrov M, Abagyan R. ICFF: A new method to incorporate implicit flexibility into an internal coordinate force field. Journal of computational chemistry. 2003;24:254-65.
[5] Schapira M, Totrov M, Abagyan R. Prediction of the binding energy for small molecules, peptides and proteins. Journal of Molecular Recognition. 1999;12:177-90.
[6] Neves MA, Totrov M, Abagyan R. Docking and scoring with ICM: the benchmarking results and strategies for improvement. Journal of computer-aided molecular design. 2012;26:675-86.
[7] Abagyan R, Totrov M. Biased probability Monte Carlo conformational searches and electrostatic calculations for peptides and proteins. Journal of molecular biology. 1994;235:983-1002.
[8] Wu X-H, Quan J-M, Wu Y-D. Theoretical study of the catalytic mechanism and metal-ion dependence of peptide deformylase. The Journal of Physical Chemistry B. 2007;111:6236-44.
